# A key regulator with a MIF-like domain orchestrates cellular differentiation and virulence in the fungal pathogen *Magnaporthe oryzae*

**DOI:** 10.1101/2022.08.05.502968

**Authors:** Matteo Galli, Stefan Jacob, Ying Zheng, Parviz Ghezellou, Martin Gand, Wendell Albuquerque, Jafargholi Imani, Valérie Allasia, Christine Coustau, Harald Keller, Bernhard Spengler, Eckhard Thines, Karl-Heinz Kogel

## Abstract

MACROPHAGE MIGRATION INHIBITORY FACTOR (MIF) is a pleiotropic protein with chemotactic, pro-inflammatory, and growth-promoting activities first discovered in mammals. In parasites, MIF homologs are involved in immune evasion and pathogenesis. Here, we present the first comprehensive analysis of a MIF protein from the devastating plant pathogen *Magnaporthe oryzae* (*Mo*). The fungal genome encodes a single MIF protein (*Mo*MIF1) that, unlike the human homolog, harbors multiple low-complexity regions (LCRs) and is unique to Ascomycota. Following infection, *MoMIF1* is expressed in the biotrophic phase of the fungus, and is strongly down-regulated during subsequent necrotrophic growth in leaves and roots. We show that *Mo*MIF1 is secreted during plant infection, affects the production of the mycotoxin tenuazonic acid and inhibits plant cell death. Our results show that *Mo*MIF1 is a novel key regulator of fungal virulence that maintains the balance between biotrophy and necrotrophy during the different phases of fungal infection.

## Introduction

In mammals and plants, cytokines are fundamental mediators of innate immunity and cell signaling (Calandra and Roger, 2003; Choi et al., 2010; Merk et al., 2012, Panstruga et al., 2015; Gruner et al., 2021). They are released after an initial binding of small molecular motifs, such as pathogen-associated molecular patterns (PAMPs), to toll-like receptors (TLRs) or pattern recognition receptors (PRRs) (Akira and Hemmi, 2003; Ausubel, 2005). MACROPHAGE MIGRATION INHIBITORY FACTOR (MIF) was one of the first cytokines discovered in humans (*Homo sapiens; Hs*). Originally, it was related to the delayed-type hypersensitivity reaction, a T-cell-mediated immune response coupled by macrophage activation and infiltration in the inflamed tissue (Bloom and Bennett, 1966; David, 1966). Since then, *Hs*MIF has been associated with a wide range of biological functions, including: *i*. production of other inflammatory cytokines (such as tumor necrosis factor, interleukin-6, interferon-γ, and interleukin-1β), *ii*. hormone immunomodulation, *iii*. regulation of muscle glucose catabolism and other cell metabolic processes, *iv*. tumor growth promotion, *v*. pivotal mediation of immune diseases such as rheumatoid arthritis and atherosclerosis, and *vi*. programmed cell death (Takahashi et al., 1998; Calandra and Roger, 2003; Fingerle-Rowson et al., 2003; Onodera et al., 2004; Morand et al., 2006; Bifulco et al., 2008; Wang et al., 2016; Nobre et al., 2017). In the human genome, *HsMIF* is located on chromosome 22 and encodes a protein of 114 amino acids (aa). Despite its limited molecular mass, the protein harbors three sequence motifs that confer distinct functional catalytic activities. These include keto-enol tautomerase, thiol-protein oxidoreductase, and DNA nuclease activities (Rosengren et al., 1997; Kleemann et al., 1998; Wang et al., 2016). *Hs*MIF also acts as a ligand for cell surface receptors such as CD74 and CXCR, thereby triggering multiple signaling pathways underlying the protein’s biological functions (Shi et al., 2006; Tillmann et al., 2013).

A considerable number of *MIF* sequences have been identified in a vast range of organisms across the eukaryotic kingdoms (Michelet et al., 2019), including nematodes, aphids, protozoans, and mollusks. The genome of the plant model *Arabidopsis thaliana* codes for three distinct MIFs (MIF/D-DT-like proteins, MDLs) and their function has been linked with plant development and modulation of innate immunity (Gruner et al., 2021). According to current knowledge, MIF in free-living organisms is mainly involved in the activation of immune responses in these species, the proliferation of immune cells and the inhibition of apoptosis mediated by p53 (Jung et al., 2008 Baeza-Garcia et al., 2010; Brock et al., 2014). However, parasitic species have been shown to produce several MIF proteins, some of which are designated for secretion and to modulate the immune responses in the host organism thus facilitating infection (Pastrana et al., 1998; Augustijn et al., 2007; Cho et al., 2011; Miller et al., 2012; Ajonina-Ekoti et al., 2013; Naessens et al., 2015; Zhao et al., 2019). Studies on the mode of action and function of such MIF proteins have mainly been conducted on human and plant parasites such as nematodes, protozoa or aphids (Augustijn et al., 2007; Vermeire et al., 2008, Naessens et al., 2015; Liu et al., 2021).

Compared to other kingdoms, very little is known about MIF proteins in fungi. Only recently, a comprehensive *in-silico* study about *MIF* evolutionary history in eukaryotes mentioned the fungal MIF protein for the first time (Michelet et al., 2019). This study showed that fungi possess no or at most one *MIF* sequence in their genome and that the predicted MIF proteins are often associated with other domains. Fungal MIFs therefore appear to be rather unusual cytokines.

To elucidate the role of MIF proteins in fungal infection strategies, we studied the filamentous plant pathogen *Magnaporthe oryzae* (*Mo*), the causal agent of devastating blast diseases on many grass species, including the cereals rice, wheat, and barley (Urashima et al., 2004; Ebbole, 2007; Faivre-Rampant et al., 2008; Cruz et al., 2012). *Mo* ranks first on the top 10 list of plant pathogenic fungi worldwide (Dean et al., 2012). This ascomycete has become a model system for fungi with broad host ranges (Routledge et al., 2004; Parker et al., 2008), and for studying the molecular basis of infection-related differentiation processes, including the lifestyle switch from biotrophic to necrotrophic growth that characterizes hemi-biotrophic fungi (Yan et al., 2016; Fernandez and Orth, 2018). *Mo* invades plant cells using intracellular invasive hyphae (IHs) that grow from one cell to the next. After penetrating the leaf cuticle from an appressorium, the IH breaches the epidermal cell wall, invaginates the plant plasma membrane, and colonizes the first-invaded cell for 8 to 12 h (Martin-Urdiroz et al., 2016; Zhu et al., 2021). In this early biotrophic stage of infection, the living host cells are metabolically reprogrammed by fungal effector proteins (Giraldo et al., 2013). Within approximately 48 hours post inoculation (hpi), secondary IHs spread into neighboring mesophyll cells, where they are enclosed by a plant membrane, called the extra-invasive hyphal membrane (EIHM). Subsequently, the fungus grows filamentously, and each adjacent plant cell is infected where the cell walls are thinnest (pit field). Newly invaded cells seem to lose their viability when the fungus moves into adjacent ones (Kankanala et al., 2007; Wilson and Talbot, 2009).

Of note, fungi of the *Magnaporthe* genus are able to penetrate and colonize cereal roots, resulting in root necrotization. A thorough analysis of the infection strategy of *M. grisea* strain Guy11 showed that melanized appressoria associated with classical foliar infection are not required for root infection (Sesma and Osbourn, 2004; Tucker et al., 2010). In contrast, hyphal swellings resembling the simple penetration structures (hyphopodia) of root-infecting fungi were visible at the infection sites. Similar to the early biotrophic growth phase in leaves, the intracellular hyphae are thick and bulbous with constrictions where they cross the plant cell.

Here we report the first comprehensive analysis of a fungal MIF protein and describe its key function in fungal virulence. We show that the genome of *Magnaporthe oryzae* harbors one *MIF* gene (*MoMIF1*), which encodes a protein that is unusual in size and structure and differs greatly from MIF homologs in all other taxa, including the closely related Basidiomycota. Using the grass model *Brachypodium distachyon* as host (Scholthof et al., 2018; Zanini et al., 2021), we discovered that *MoMIF1* is a highly regulated fungal gene that is transiently up-regulated in the very early biotrophic phase but strongly down-regulated in the subsequent necrotrophic phase of plant colonization. Analyses of *MoMIF1* knock-out (KO) and overexpressor (OE) mutants confirmed that *Mo*MIF is detrimental to the necrotrophic phase of fungal development in leaves and roots. Consistent with this, we show that *Mo*MIF1 is a secreted protein with cell death inhibitory activity. Our results suggest that *Mo*MIF1 determines fungal development and virulence by regulating the pathogen’s lifestyle switch during infection.

## Results

### Ascomycota MIFs have unique features

Using the *HsMIF* gene sequence as reference, we conducted a genome-wide comparison of *MIF* sequences from the EnsemblFungi database (Howe et al., 2020). In the subkingdom Dikarya, where most of the plant pathogenic fungi are listed, we found one *MIF*-like gene per genome (Table 1), supporting our previous data on a broad survey of MIF sequences between different kingdoms (Michelet et al., 2019). Intriguingly, Ascomycota MIFs are much larger proteins (~3 times the size of the *Hs*MIF), compared with MIFs from model organisms of different kingdoms, including Animalia, Plantae, Protozoa and to other Fungi (Fig. 1a). The tautomerase/MIF domain motif is located roughly in the middle of the sequence, flanked by long peptide “arms”, which carry multiple low complexity regions (LCRs) of unknown function. Phylogenetic analysis with predicted MIF domain sequences from plants, animals, invertebrates and fungi confirmed that Ascomycota MIFs form a separate clade (Fig. S1); taking the *Mo*MIF1 domain (MGG_05693) as input sequence, we calculated Ascomycota MIF sequence similarity ranging from 59% to 88%, while Basidiomycota and other taxa MIF domain sequences scored lower (between 50% and 19%; Table 1). Multiple sequence alignment analyses revealed that *Mo*MIF1 lacks CxxCxxHx(n)C (red underlined) and PD-D/E(x)K (green highlighted) motifs, thus suggesting absence of oxidoreductase and DNA nuclease activities, respectively. Furthermore, the active site residues for the tautomerase activity (white triangles) are not conserved (Fig. 1b).

**Fig. 1.**
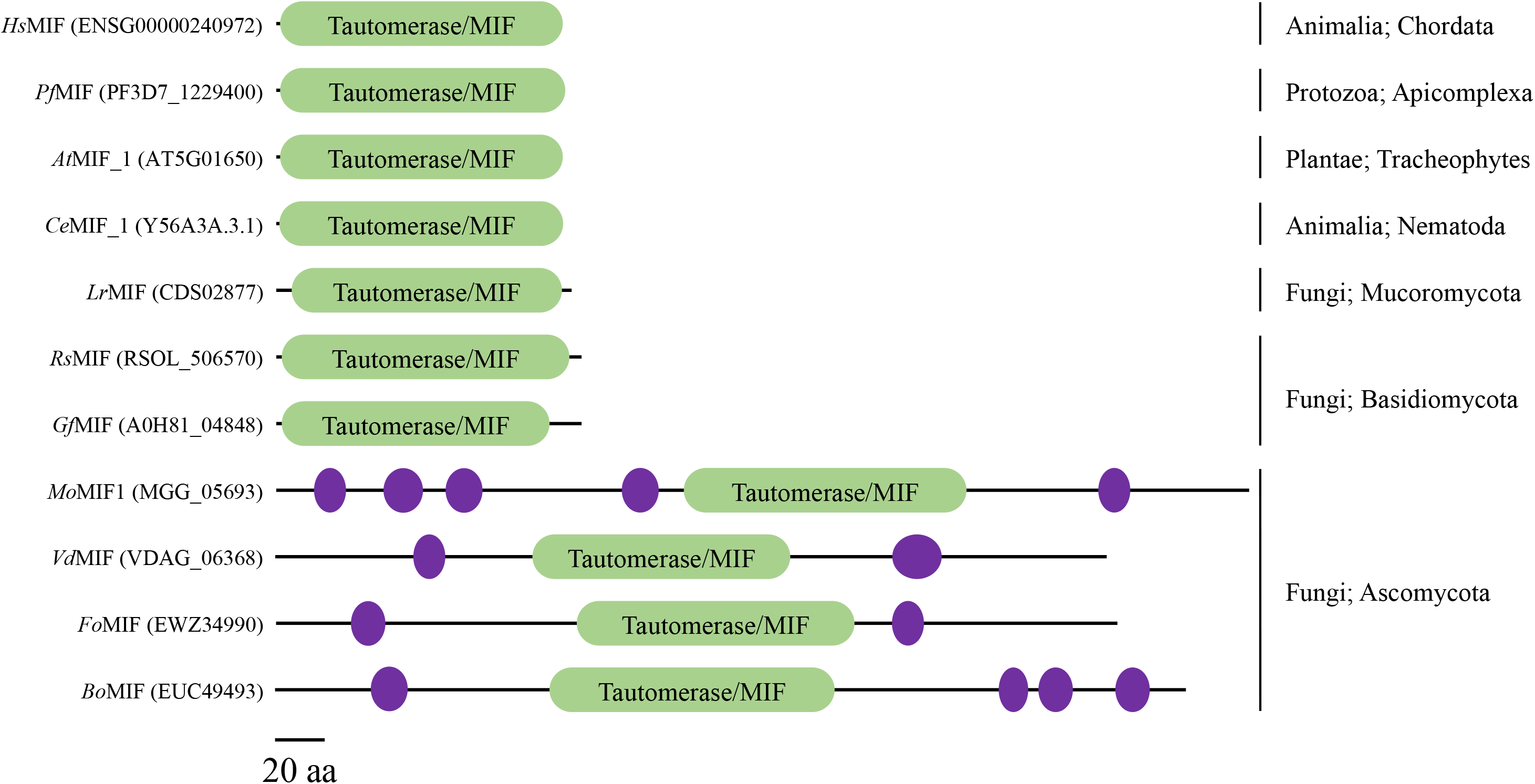
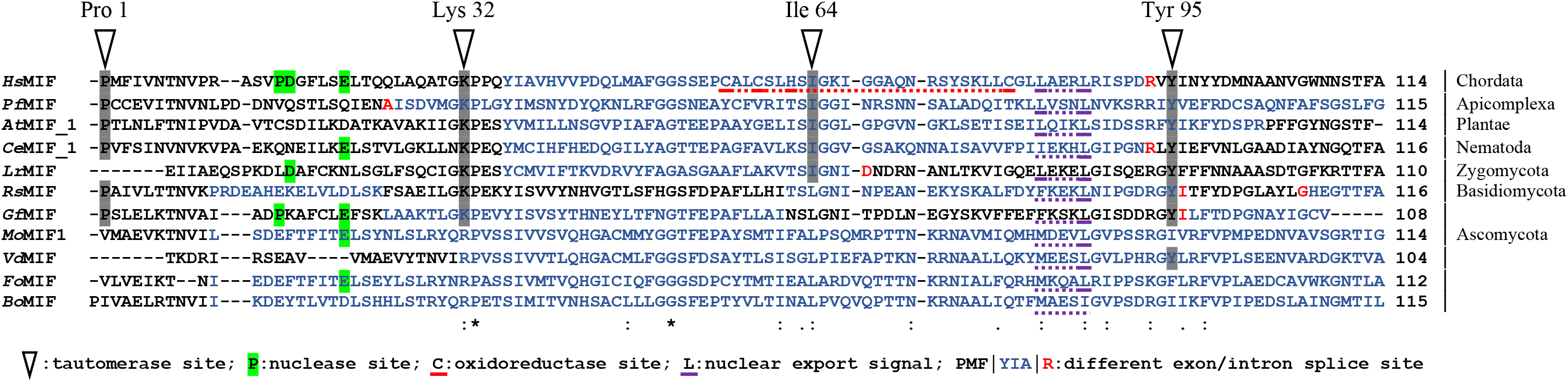
MIF protein domain and structure comparison. (a) Protein organization of *Magnaporthe oryzae* (*Mo*MIF1, MGG_05693) with its homologs in *Homo sapiens* (*Hs*), the protozoan parasite *Plasmodium falciparum* (*Pf*), the genetic plant model organisms *Arabidopsis thaliana* (*At*) and *Caenorhabditis elegans* (*Ce*), one parasitic Mucoromycota fungus *Lichtheimia ramosa* (*Lr*), two plant pathogenic Basidiomycota fungi *Rhizoctonia solani* (*Rs*) and *Grifola frondosa* (*Gf*) and three plant pathogenic Ascomycota fungi *Verticillium dahlia* (*Vd*), *Fusarium oxysporum* (*Fo*) and *Bipolaris oryzae* (*Bo*). Domain structure comparison via the online-tool CDD/SPARCLE (Marchler-Bauer et al., 2017). Green marker highlights the Tautomerase/MIF domain, black line represents length of the protein (in scale), and purple circle denotes low complexity region of unknown function. (b) Conservation of amino acids (aa) sites and motifs in consensus MIF sequences. Hyphens (-) indicate a gap in the respective sequence. Alternance of black and blue color indicates difference exons, red aa indicate that the residue overlaps splice site. Aa sequences marked by symbols (above or below) define distinct functional catalytic activities in human MIF. Tautomerase activity (triangle symbol), oxidoreductase activity (CxxCxxHx(n) motif; red underlined), nuclease activity (PE/DxxxxE motif; green highlighted) and nuclear export signal (LxxxL; purple underlined). Conserved aa residues between species in the tautomerase domain are grey highlighted. Below the protein sequences is a key denoting conserved sequence (*), conservative mutations (:), and semi-conservative mutations (.).

**Table 1.** Comaparison of the *Mo*MIF1 domain with various eukaryotic MIF domains as identified by species, accession numbers, total protein length, position of the MIF domain in the protein, confidently E-value of the domain and aa similarity to *Mo*MIF1 domain. Domain structure comparison via the online-tool SMART with PFAM (Letunic et al., 2021). Similarity analysis via EMBOSS needle (Needleman and Wunsch, 1970). *: fungus, Ascomycota; °: fungus, non-Ascomycota (Basidiomycota, Oomycota, Mucoromycota and Glomeromycota); ^: other non-fungal organism; aa: amino acid; bp: base pair.

| Species | Accession number [Ensembl] | Protein length [aa] | MIF domain position [bp] | MIF domain E-value | MIF domain similarity to MoMIF1 |
| --- | --- | --- | --- | --- | --- |
| *Magnaporthe oryzae* * | MGG_05693T0 | 392 | 164 to 277 | 2.80E-11 | / |
| *Podospora anserina* * | CAP68549 | 334 | 84 to 197 | 1.20E-12 | 87.70% |
| *Neurospora crassa* * | KHE87850 | 485 | 171 to 284 | 7.10E-11 | 84.20% |
| *Bipolaris oryzae* * | EUC49493 | 366 | 111 to 225 | 4.90E-16 | 82.60% |
| *Colletotrichum graminicola* * | GLRG_03638 | 373 | 118 to 232 | 2.10E-14 | 81.70% |
| *Erysiphe necator* * | KHJ30970 | 335 | 118 to 231 | 6.60E-12 | 78.90% |
| *Verticillium longisporum* * | CRK39038 | 692 | 114 to 226 | 3.00E-13 | 78.90% |
| *Microdochium bolleyi* * | KXJ97725 | 392 | 160 to 273 | 3.00E-11 | 78.90% |
| *Zymoseptoria tritici* * | SMQ48239 | 724 | 125 to 238 | 2.80E-16 | 77.20% |
| *Blumeria graminis f. sp. Tritici* * | EPQ64960 | 338 | 121 to 234 | 1.10E-10 | 77.20% |
| *Rhynchosporium commune* * | CZS95638 | 338 | 126 to 238 | 7.00E-11 | 75.40% |
| *Rhynchosporium secalis* * | CZT52938 | 338 | 126 to 238 | 5.10E-11 | 75.40% |
| *Fusarium graminearum* * | ESU11401 | 343 | 120 to 233 | 4.90E-10 | 72.80% |
| *Botrytis cinerea* * | Bcin10g02580.1 | 348 | 138 to 252 | 1.50E-12 | 72.20% |
| *Sclerotinia sclerotiorum* * | APA10581 | 353 | 139 to 253 | 6.60E-12 | 72.20% |
| *Fusarium culmorum* * | FCUL_07598.1 | 343 | 121 to 233 | 2.30E-06 | 71.10% |
| *Trichoderma harzianum* * | KKO98006 | 325 | 106 to 222 | 5.00E-12 | 70.90% |
| *Fusarium oxysporum* * | EWZ34990 | 338 | 122 to 233 | 1.40E-12 | 69.30% |
| *Sclerotinia borealis* * | ESZ93718 | 336 | 130 to 236 | 1.20E-11 | 64.30% |
| *Aspergillus niger* * | SPB43847 | 351 | 137 to 250 | 1.00E-09 | 64.00% |
| *Tuber magnatum* * | PWW80050 | 330 | 114 to 226 | 5.60E-15 | 62.30% |
| *Tuber borchii* * | PUU79916 | 330 | 114 to 226 | 3.60E-15 | 62.30% |
| *Tuber melanosporum* * | CAZ84338 | 331 | 114 to 226 | 3.60E-17 | 61.40% |
| *Penicillium digitatum* * | EKV13723 | 369 | 121 to 235 | 4.40E-10 | 60.90% |
| *Fonsecaea pedrosoi* * | KIW84848 | 326 | 124 to 236 | 2.40E-13 | 60.50% |
| *Verticillium dahliae* * | EGY14878 | 335 | 103 to 206 | 1.00E-11 | 60.50% |
| *Exophiala aquamarina* * | KEF57016 | 334 | 131 to 242 | 1.20E-12 | 59.60% |
| *Aspergillus oryzae* * | OOO12748 | 355 | 143 to 250 | 1.90E-07 | 58.80% |
| *Verticillium alfalfae* * | EEY22597 | 340 | 99 to 206 | 1.90E-12 | 58.60% |
| *Phellinus noxius*° | PAV19534 | 116 | 2 to 114 | 1.40E-23 | 50.00% |
| *Punctularia strigosozonata*° | EIN04787 | 119 | 2 to 114 | 1.30E-19 | 49.60% |
| *Laccaria bicolor*<sup>°</sup> | EDR10872 | 120 | 2 to 114 | 1.40E-24 | 49.10% |
| *Pleurotus ostreatus*<sup>°</sup> | KDQ33921 | 120 | 2 to 114 | 3.90E-25 | 48.20% |
| *Fistulina hepatica*<sup>°</sup> | KIY45470 | 118 | 2 to 114 | 3.30E-24 | 48.20% |
| *Coprinopsis cinerea*<sup>°</sup> | EAU88060 | 124 | 2 to 119 | 9.80E-21 | 45.40% |
| *Daedalea quercina*<sup>°</sup> | KZT73998 | 119 | 2 to 114 | 8.00E-21 | 45.10% |
| *Arabidopsis thaliana* #2<sup>^</sup> | AT5G01650.1 | 115 | 2 to 115 | 2.60E-25 | 44.80% |
| *Caenorhabditis elegans* #2<sup>^</sup> | C52E4.2a.1 | 120 | 2 to 115 | 5.10E-52 | 44.50% |
| *Caenorhabditis elegans* #3<sup>^</sup> | F13G3.9.1 | 146 | 2 to 113 | 1.70E-23 | 44.30% |
| *Galerina marginata*<sup>°</sup> | KDR81866 | 120 | 2 to 114 | 3.60E-21 | 43.00% |
| *Rhizopus microsporus*<sup>°</sup> | PHZ17589 | 122 | 2 to 114 | 5.30E-21 | 43.00% |
| *Homo sapiens*<sup>^</sup> | ENST00000215754.8 | 115 | 2 to 115 | 6.40E-55 | 42.60% |
| *Phytophthora infestans* #1<sup>°</sup> | PITG_16205T0 | 116 | 2 to 116 | 6.40E-28 | 42.00% |
| *Rhizoctonia solani*<sup>°</sup> | EUC66088 | 123 | 2 to 117 | 1.20E-24 | 40.80% |
| *Plasmodium falciparum*<sup>^</sup> | CZT99452 | 116 | 2 to 116 | 3.10E-20 | 40.50% |
| *Physcomitrium patens*<sup>^</sup> | Pp3c13_4090V3.1 | 115 | 2 to 115 | 2.00E-33 | 40.50% |
| *Grifola frondosa*<sup>°</sup> | OBZ75583 | 123 | 2 to 109 | 1.00E-22 | 40.50% |
| *Phytophthora infestans* #2<sup>°</sup> | PITG_16206T0 | 120 | 2 to 115 | 7.30E-25 | 40.30% |
| *Caenorhabditis elegans* #1<sup>^</sup> | Y56A3A.3.1 | 117 | 2 to 117 | 1.10E-28 | 40.30% |
| *Caenorhabditis elegans* #4<sup>^</sup> | Y73B6BL.13.1 | 121 | 2 to 117 | 2.90E-15 | 39.80% |
| *Mucor ambiguus*<sup>°</sup> | GAN02096 | 118 | 2 to 114 | 4.00E-18 | 39.30% |
| *Arabidopsis thaliana* #1<sup>^</sup> | AT5G57170.1 | 115 | 2 to 115 | 1.40E-25 | 37.40% |
| *Serendipita indica* #1<sup>°</sup> | CCA75852 | 361 | 9 to 90 | 1.90E-08 | 37.10% |
| *Arabidopsis thaliana* #3<sup>^</sup> | AT3G51660.1 | 112 | 2 to 110 | 7.10E-13 | 35.40% |
| *Lichtheimia corymbifera*<sup>°</sup> | CDH53730 | 119 | 4 to 114 | 2.10E-15 | 34.20% |
| *Lichtheimia ramosa*<sup>°</sup> | CDS02877 | 119 | 5 to 114 | 2.40E-13 | 34.10% |
| *Serendipita indica* #2<sup>°</sup> | CCA75762 | 79 | 1 to 77 | 1.50E-12 | 28.00% |
| *Rhizophagus irregularis*<sup>°</sup> | PKK71463 | 88 | 34 to 88 | 0.0000005 | 19.30% |

### MoMIF1 expression is transiently upregulated during fungal germination and penetration

To get a first indication about the requirement of *Mo*MIF for fungal infection, *MoMIF1* expression was recorded in the time range of 0 to 96 hpi, which covers the biotrophic and the early necrotrophic phase of plant colonization. Leaves of *Brachypodium distachyon* Bd21-3 seedlings, which are highly susceptible to *Magnaporthe oryzae* strain 70-15 (Zanini et al., 2021), were spray-inoculated with conidia and *MoMIF1* expression was measured by RT-qPCR. We detected a high expression of *MoMIF1* transcripts during the early colonization phase till 12 hpi followed by a steady decline until *MoMIF1* mRNA was no longer detectable between 36 and 42 hpi (Fig. 2a). Similarly, upon dip inoculation of roots, *MoMIF1* expression was even more transient with a slightly delayed maximum at 8 hpi, followed by a strong decline in transcript abundance as early as 12-16 hpi (Fig. 2b). To further specify the exact timing of accumulation and localization of *Mo*MIF1, we generated a *Mo* reporter strain Δ*Mo*MIF1+GFP that contained a chimeric *MIF1-GFP* gene under the control of the native *MIF1* promoter. Using confocal laser scanning microscopy (CLSM), we examined the pre-infection phase starting with conidia germination on plastic hydrophobic coverslips that are known to induce appressorium formation in *Magnaporthe* (Lee and Dean, 1994; Thines et al., 2000). Prior to germination, GFP fluorescence was found exclusively in the proximal cell of the three-celled conidia from which the germ tube protrudes, confirming that *MoMIF1* is already expressed in non-germinated conidia. Further analyses over a period of 12 h after conidial germination confirmed accumulation of MIF1-associated fluorescence in both germ tubes and appressoria (Fig. 2c). In contrast, *MoMIF1* was not expressed in mycelia from a 5-day-old liquid cultures (Fig. 2d). The above data are consistent with the hypothesis that *MoMIF1* is differentially regulated in the bio- and necrotrophic colonization phases and is thus required for successful infection by the hemi-biotrophic fungus.

**Fig. 2.**
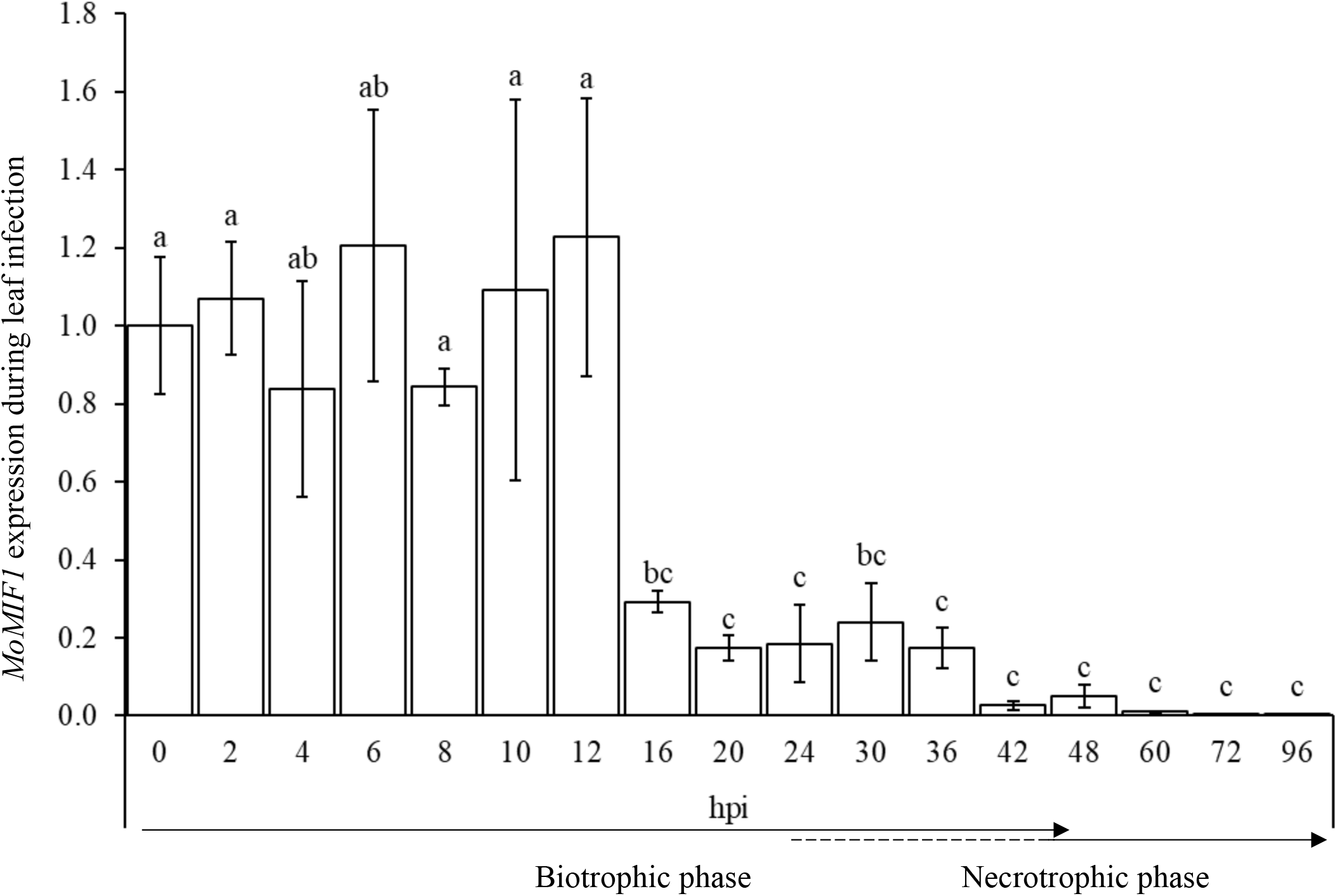
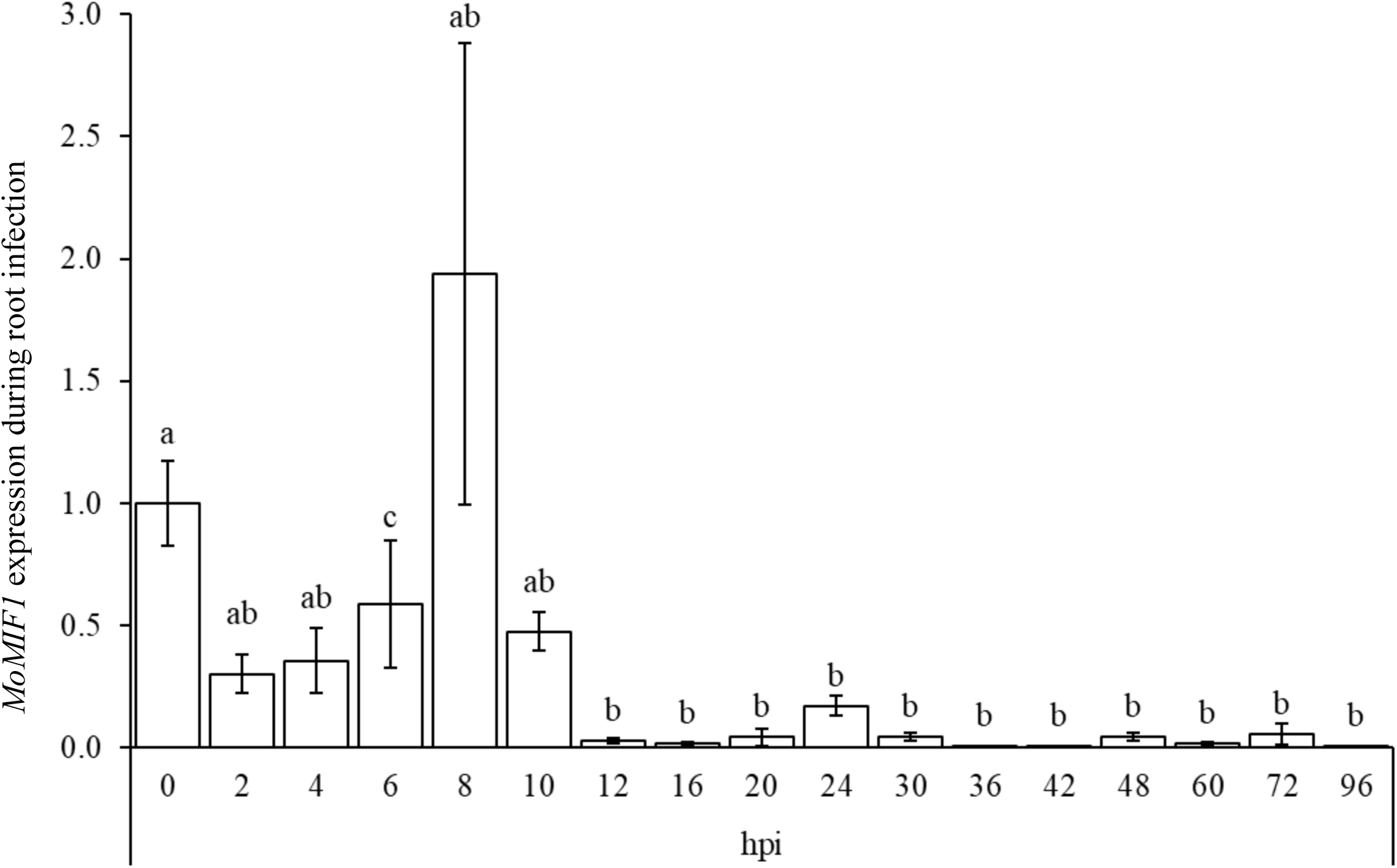
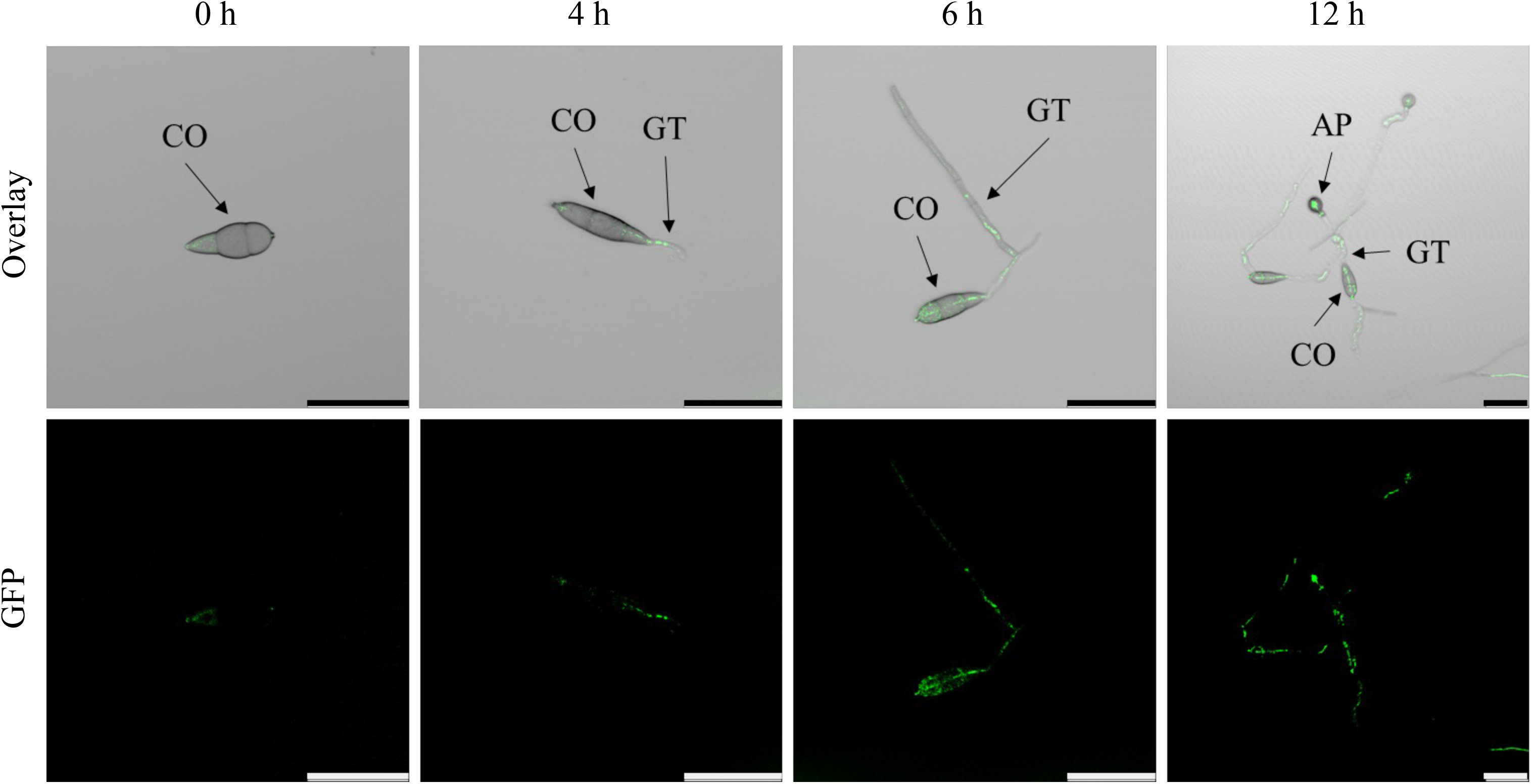
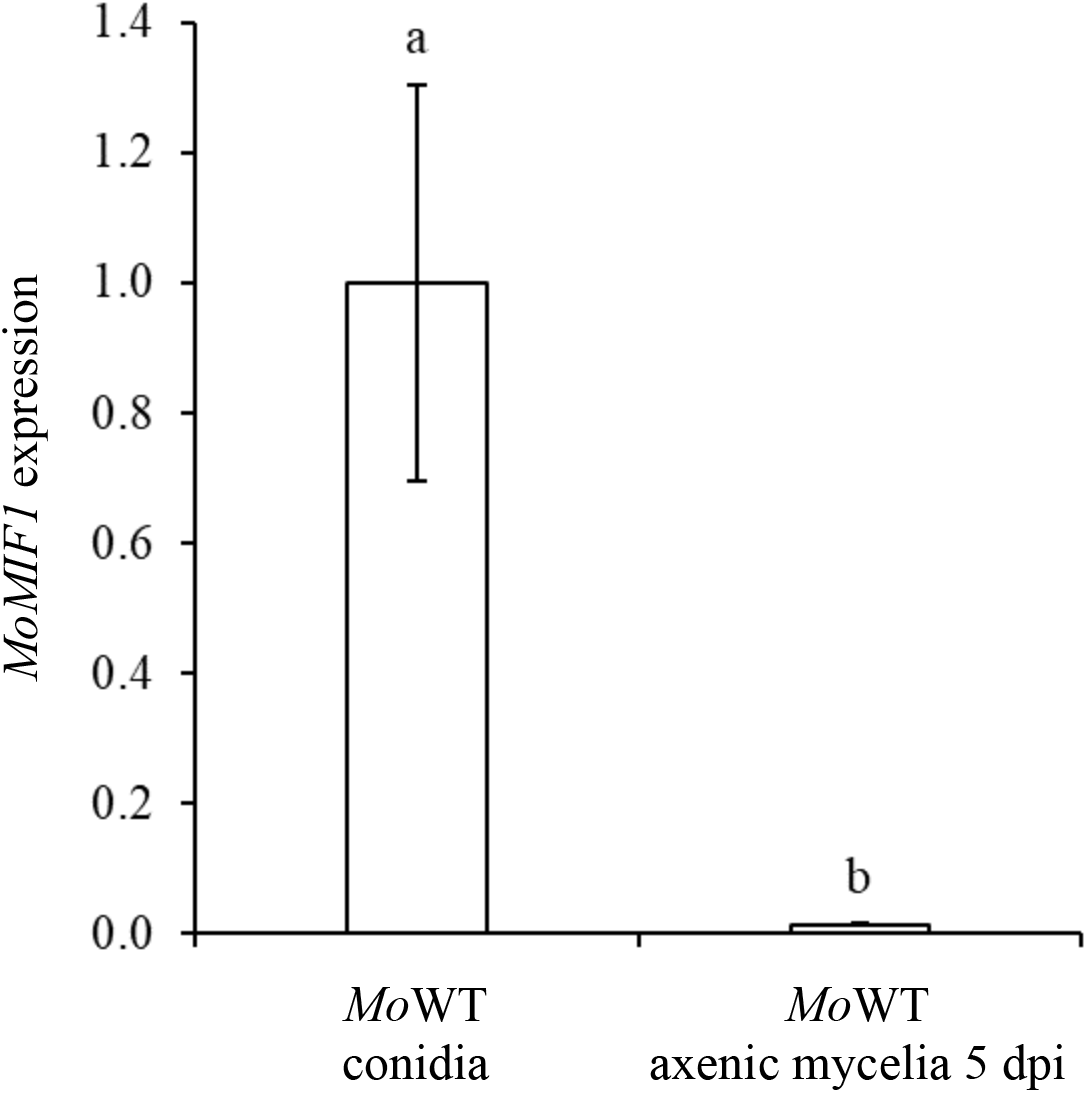
Analysis of *MoMIF1* mRNA and protein localization during *M. oryzae* development. Relative *MoMIF1* expression in WT strain 70-15 during fungal development on leaves (a) and roots (b) of the grass model plant *Brachypodium* Bd21-3 as measured by RT-qPCR. Samples were taken from the second youngest spray-inoculated leaf of whole plants (n=8) and from dip-inoculated roots (n=6) until 96 hpi, respectively. Inoculum concentration was 120 x 10^3^ conidia ml^-1^ in 0.002% Tween20 water. Fungal *glyceraldehyde-3-phosphate dehydrogenase (MoGPD*) was used for normalization. Shown is the mean (±SE) from three biological replicates. Comparisons between groups were performed via ANOVA and pairwise t-tests with Bonferroni’s correction. Letters represent statistical differences among all group means (α < 0.05). In the leaf infection graph, time course of the developmental switch during pathogenesis by *Mo* are taken form Parker et al. (2008). The 0 hpi *MoMIF1* expression sample is shared in leaf and root infection. (c) GFP signals of *Mo*MIF1 C-terminal fused with GFP, detected in germinating Δ*Mo*MIF1+GFP macroconidia (5 x 10^3^ ml^-1^ in 0.002% Tween20) grown on hydrophobic coverslips for 12 h. Scale bar: 100 μm. CO: conidia, GT: germ tube, AP: appressoria; Overlay: GFP: green fluorescence (λ_emission_ (nm): 508; λ_excitation_ (nm): 489). (d) RT-qPCR analysis of *MoMIF1* expression in freshly harvested conidia vs. 5-day-old axenic cultures. *MoGPD* was used as normalization gene. Shown is the mean (±SD) from three biological replicates. Comparisons between groups were performed via ANOVA and pairwise t-tests with Bonferroni’s correction. Letters represent statistical differences among group means (α < 0.05).

### Both KO and OE of MoMIF1 renders M. oryzae less virulent on Bd21-3 leaves

To further explore the function of *Mo*MIF1 during plant infection, we generated knock-out (KO) mutants (Δ*Mo*MIF1-KO_1, Δ*Mo*MIF1-KO_2), overexpressor (OE) mutants (Δ*Mo*MIF1-OE_1, Δ*Mo*MIF1-OE_2), and a complementation mutant Δ*Mo*MIF1-comp (containing *MoMIF1* driven by the original *MIF1* promoter). Southern blot analysis proved the correct manipulation of the fungal genome (Fig. S2a), and RT-qPCR analysis confirmed strong *MoMIF1* expression in 5-day-old mycelia of Δ*Mo*MIF1-OE_1 and Δ*Mo*MIF1-OE_2 but not in KO mutants and wild type (WT) (Fig. S2b). The KO mutants’ germination rate, size, morphology and strength of appressoria measured by the number of appressoria that collapsed after 5 min over a wide range of glycerol concentrations was indistinguishable from the wt (Fig. S3a). Moreover, we could not detect substantial differences in mycelium pigmentation, appearance and conidiation on a range of rich (CM) and minimal media (MM) between KO mutants and WT, suggesting that *Mo*MIF1 is dispensable for saprophytic growth (Fig. S3b).

The above data together indicated a role of *Mo*MIF1 in the fungal infection process. Therefore, we assessed the virulence of KO and OE mutants on Bd21-3 leaves. Leaves were spray-inoculated and at 5 days post infection (dpi) the total size of the diamond-shaped necrotic blast lesions was recorded. KO mutants elicited significantly smaller necrotic lesions compared to WT (Fig. 3a,b), and the amount of fungal DNA extracted from infected leaves was significantly lower (Δ*Mo*MIF1-KO_1: 55%; Δ*Mo*MIF1-KO_2: 66% less DNA vs. WT; Fig. 3c). Unexpectedly, OE mutants were also strongly impaired in virulence (Δ*Mo*MIF1-OE_1: 79%; Δ*Mo*MIF1-OE_2: 89% reduction in fungal DNA vs. WT), while complementation mutant Δ*Mo*MIF1-comp almost completely restored the WT virulence phenotypes (Fig. 3a-c). Of note, and further supporting the above data, both KO and OE mutants showed also lower virulence on rice (*Oryzae sativa* cv. CO-39) leaves compared with WT (Fig. S4). Taken together, our data suggested that the MIF protein of *Mo* is required for virulence and its function is closely linked to the early infection phase, while it seems to have a negative effect on the later necrotrophic development of the fungus.

**Fig. 3.**
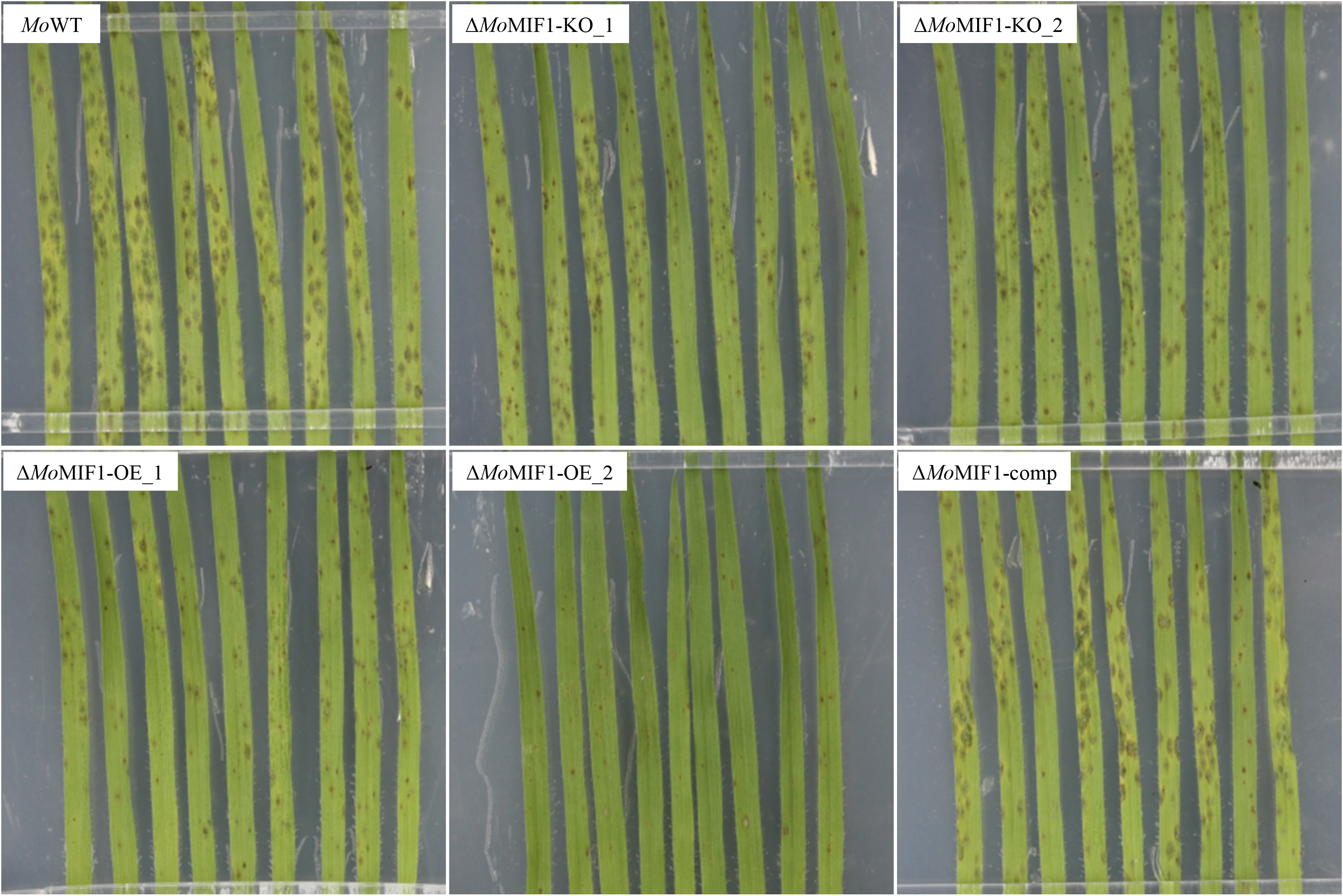
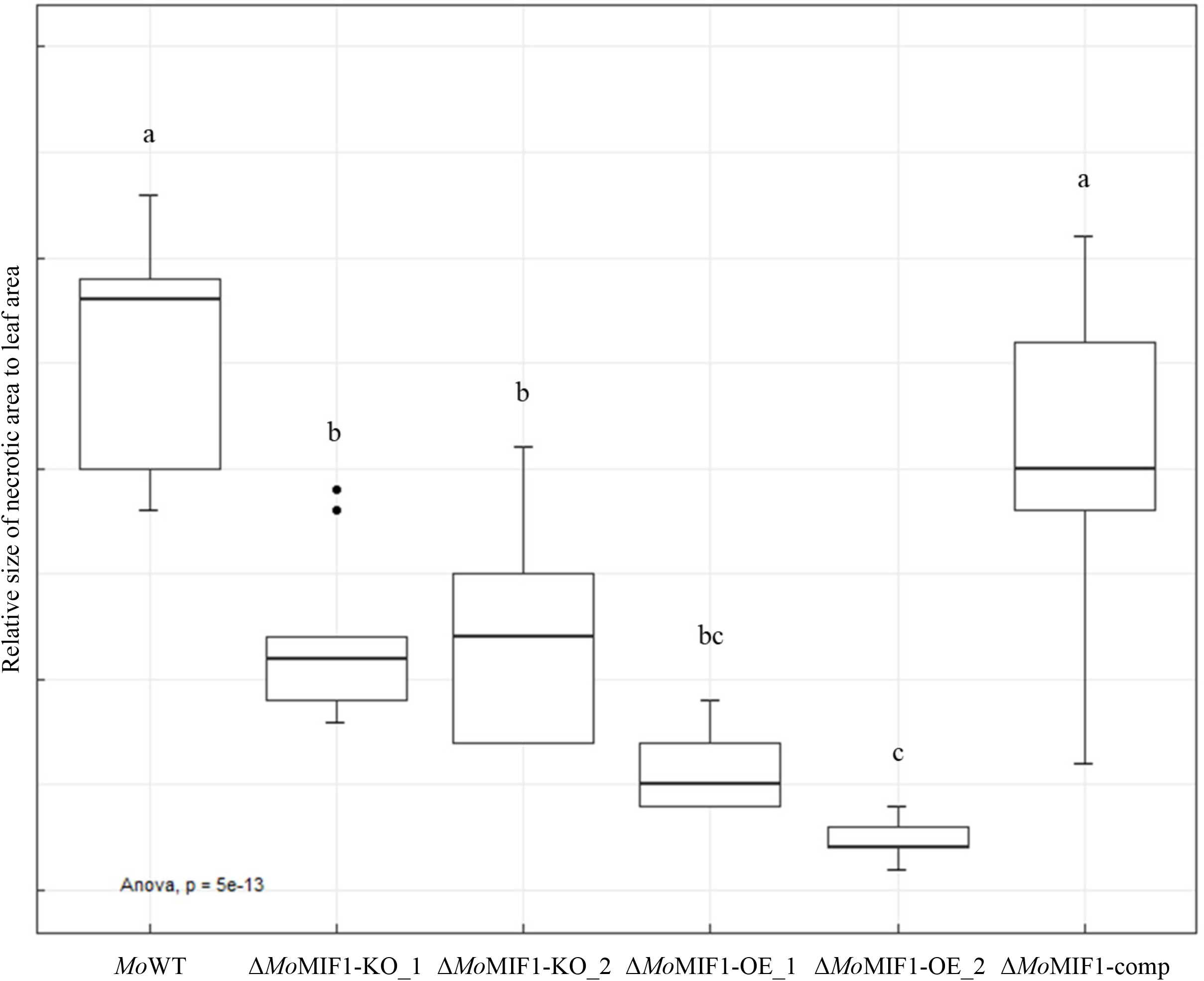
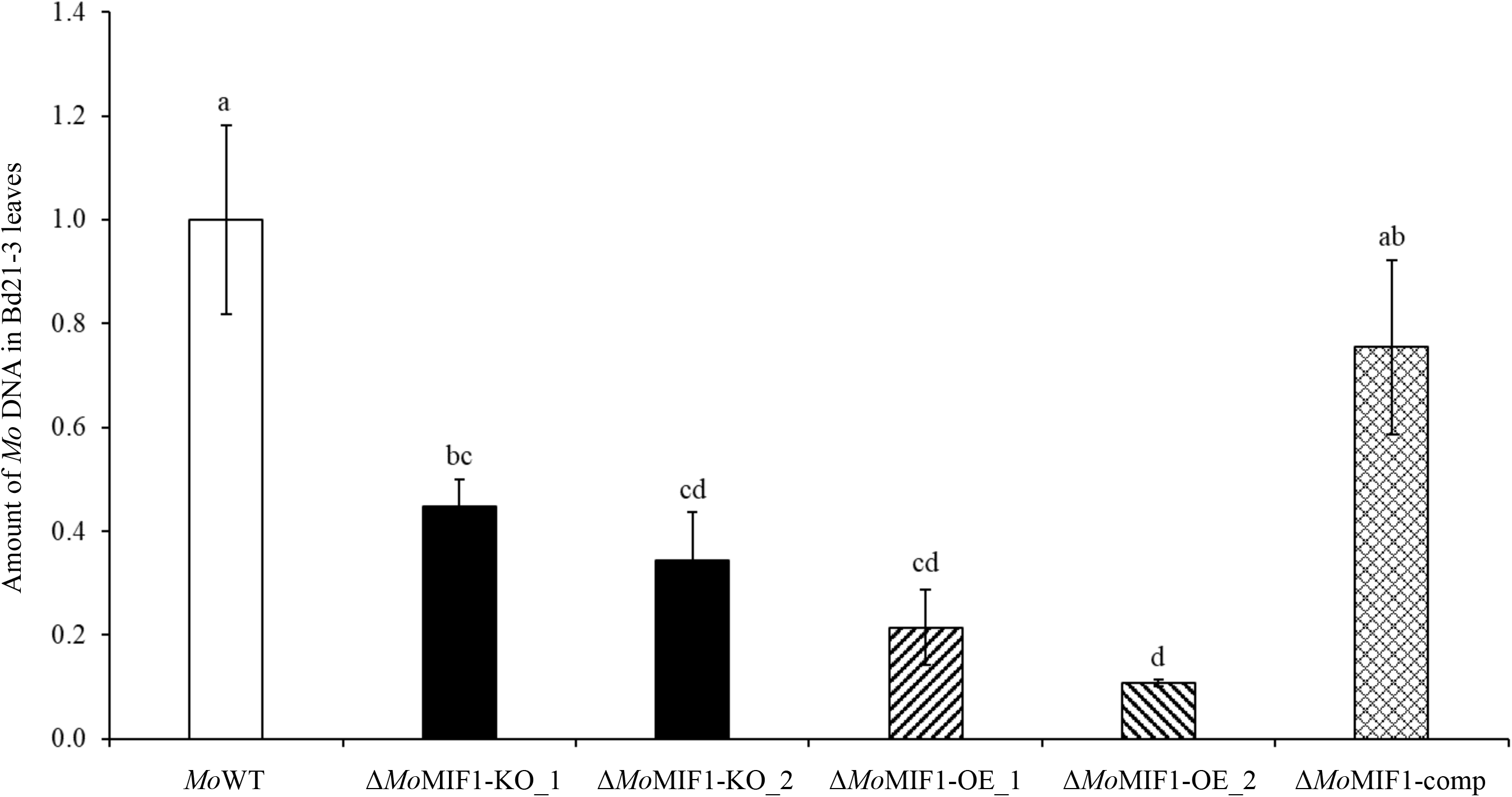
Disease symptoms caused by *Magnaporthe oryzae* 70-15 on *BIm distachyon* Bd21-3 leaves. (a) Phenotypical analysis of the infection phenotypes of WT, *MoMIF1* KO and OE mutants, and complementation mutant Δ*Mo*MIF1-comp. Three-week-old seedlings were sprayed with a suspension of 120 x 10^3^ conidia ml^-1^ in 0.002% Tween20 and kept for 5 days under high humidity at 16 h light/8 h dark cycle at 22°C/18°C. (b) Size of the necrotic area relative to total leaf size as assayed via ImageJ software at 5 dpi. (c) Quantification of fungal DNA in second youngest leaves (n=9 plants) at 5 dpi via qPCR (ratio fungal *GPD* normalized to plant *ubiquitin*). In this experiment two independent KO mutants (Δ*Mo*MIF1-KO_1, Δ*Mo*MIF1-KO_2), and two independent OE mutants (Δ*Mo*MIF1-OE_1, Δ*Mo*MIF1-OE_2) were used. Shown is the mean (±SD) from three biological replicates. Comparisons between groups were performed via ANOVA’ and Tukey’s Range Test. Letters represent statistical differences among all group means (α = 0.05).

### KO mutants, but not OE mutants heavily colonize Bd21-3 roots

The above assumption prompted us to test the virulence of mutants on Bd21-3 roots. *Mo* penetrates cereal roots without a requirement for appressoria. Tissue colonization is subsequently characterized by a very short biotrophic phase followed by extensive necrotrophic infection (Sesma and Osbourn, 2004; Marcel et al., 2010; Tucker et al., 2010). We thus performed root infection assays to determine whether *Mo*MIF1 is required for root penetration and invasion. Bd21-3 roots from five-day-old seedlings were dip-inoculated with conidia of WT and all mutants. At 5 dpi, roots were carefully removed from the substrate, cleaned, subjected to visual analysis and quantification of fungal DNA. Roots infected with Δ*Mo*MIF1-KO_1 were heavily colonized (Fig. 4), were shorter and exhibited more severe root browning compared with WT (Fig. S5). In contrast, roots inoculated with Δ*Mo*MIF1-OE_1 showed a slightly reduced necrotization and, consistent with this, colonization was slightly but not significantly reduced as evidenced by lower amounts of fungal DNA as compared with the WT, while Δ*Mo*MIF1-comp displayed similar root necrotization and infection as the WT (Fig. 4; Fig. S5). These data further substantiate that i. *Mo*MIF1 is adverse to necrotrophic growth, and that ii. *Mo*MIF1 is not as critical to early root infection as it is to early leaf infection, which is consistent with the knowledge that the biotrophic phase of root invasion is very short followed by extensive necrotrophic infection.

**Fig. 4.**
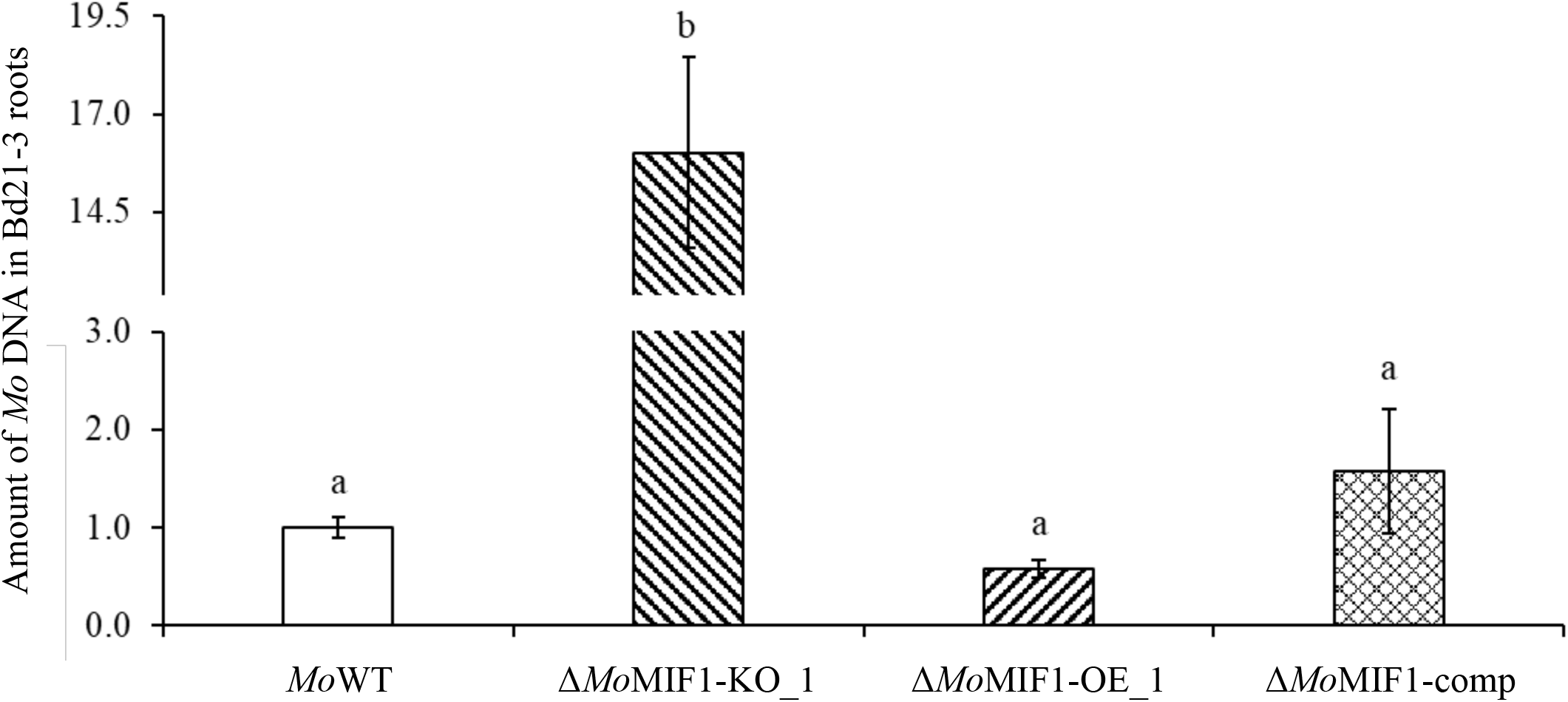
Quantification of fungal DNA in *Brachypodium distachyon* Bd21-3 roots (n=10 plants) infected with *Mo*WT and MIF mutants at 5 dpi as measured by qPCR based on the ratio of fungal *actin* vs. plant *ubiquitin*. Shown is the mean (±SD) from three biological replicates. Comparisons between’ groups were performed via ANOVA and Tukey’s Range Test. Letters represent statistical differences among all group means (α = 0.05).’

### MoMIF1 is detrimental to hyphal expansion into mesophyll cells

The above results are consistent with the idea that *Mo*MIF1 is required at very early stages of fungal infection, while it is rather harmful in later fungal development during necrotizing colonization of plant tissue. To further explore this hypothesis, we injected 100 μl of conidia solution of *MoMIF1* WT and mutants directly into the apoplastic space of second-youngest leaves of three-week-old Bd21-3 seedlings. Injection of conidia circumvents the leaf penetration stage (appressorium-mediated infection and invasion) and shortens the phase of biotrophic growth. We found that injected *MoMIF1* KO mutants were even more virulent than the WT, causing a higher number and larger blast lesions (Fig. 5a,b). In marked contrast, and consistent with our expectation, injection of *MoMIF1* OE mutants resulted in less infection and substantially reduced necrotic blast symptoms compared to KO mutants (Fig. 5a,b). Together with the data from root infections, this shows that *Mo*MIF1 is required for the early penetration stage and biotrophic leaf invasion, while it is detrimental to the necrotrophic development of the fungus.

**Fig. 5.**
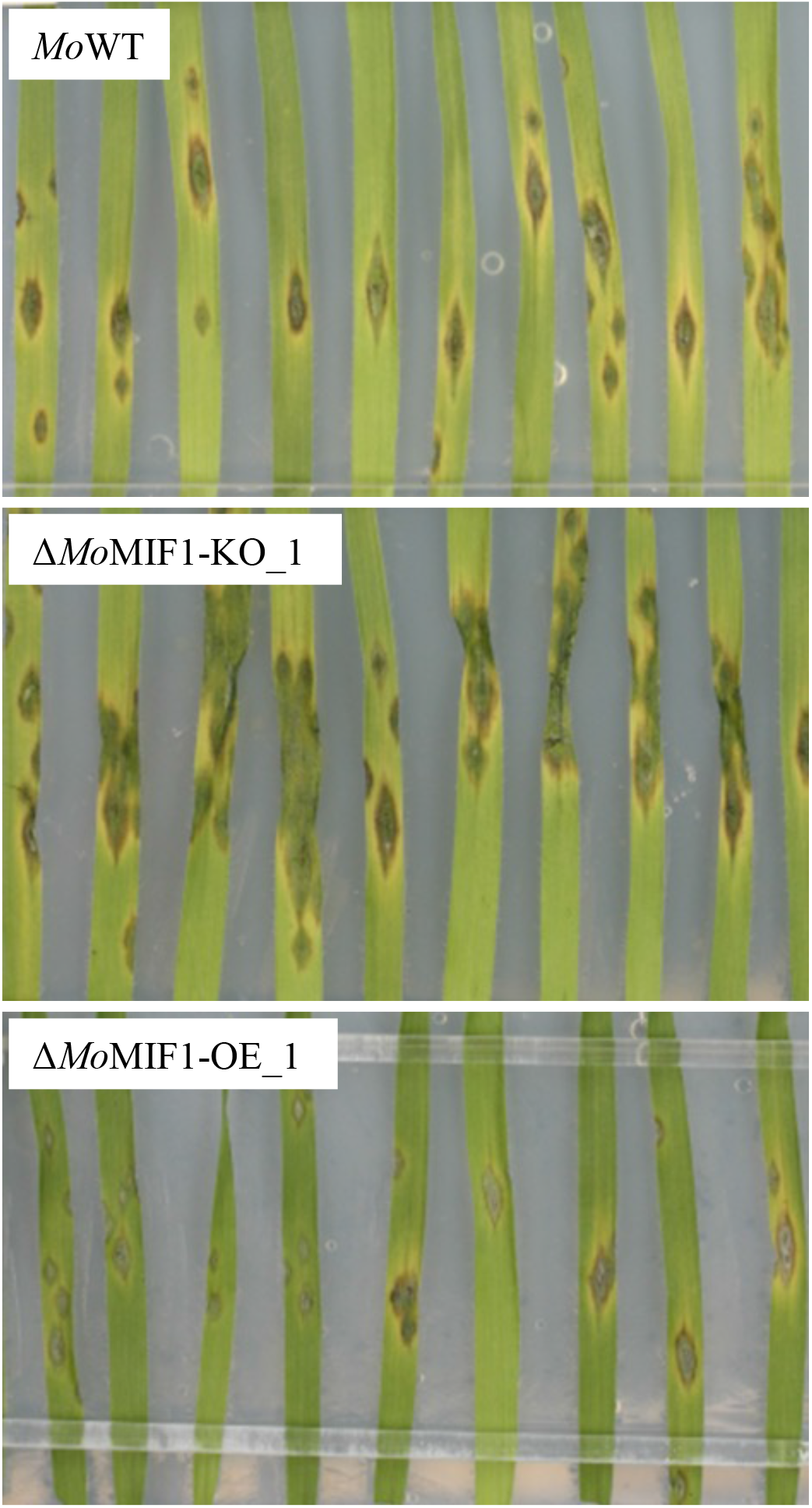
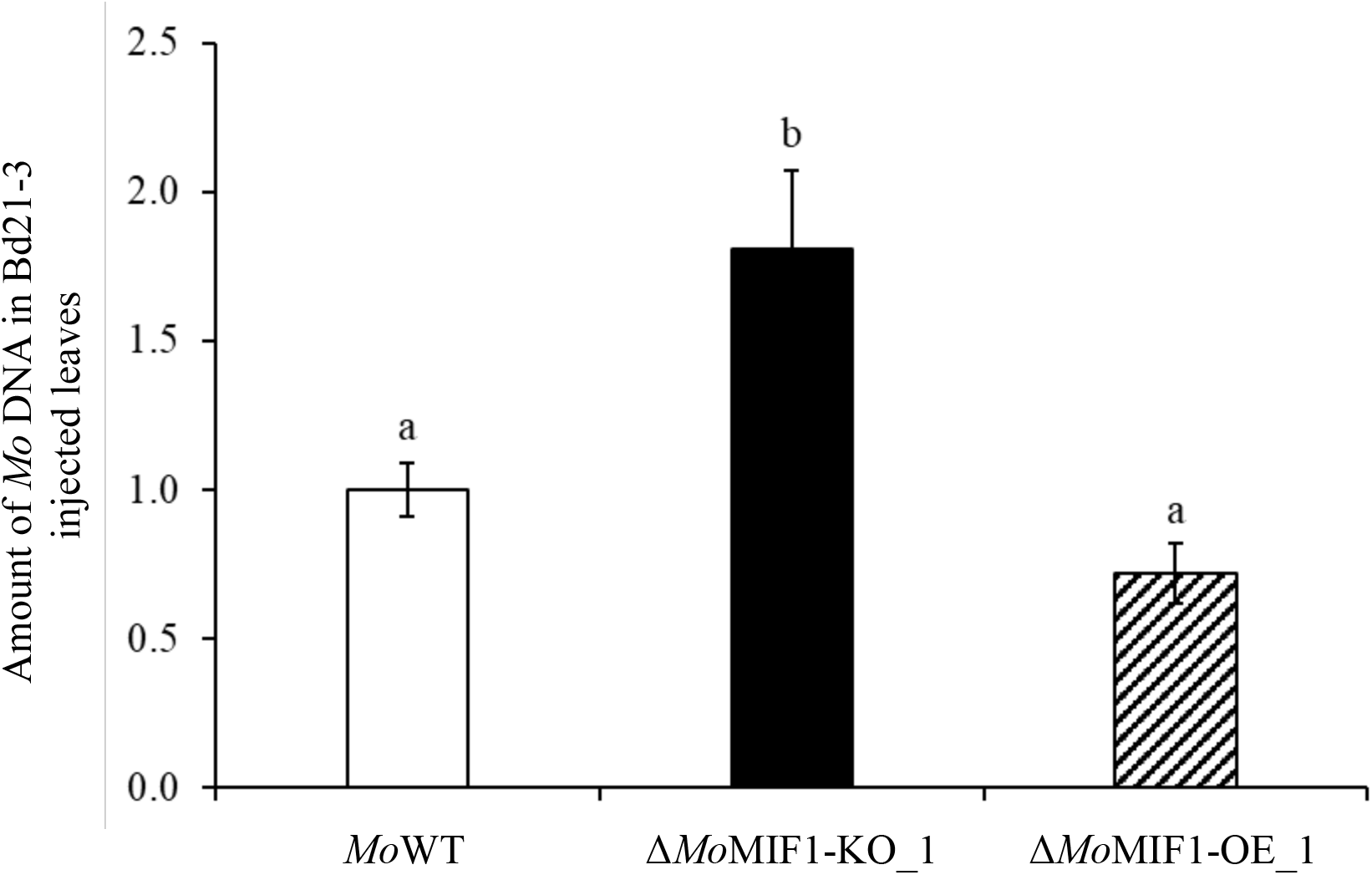
Disease symptoms caused by injection of *Mo* conidia into the apoplastic space of Bd21-3. (a) Phenotypical analysis of the injection infection phenotypes of WT, *MoMIF1* KO and OE mutants. The second youngest leaf of three-week-old seedlings (n=8 plants) were injected with a suspension of 120 x 10^3^ conidia ml^-1^ in 0.002% Tween20 and kept for 5 days under high humidity at 16 h light/8 h dark cycle at 22°C/18°C. (b) Quantification of fungal DNA at 5 dpi via qPCR (ratio fungal *GPD* normalized to plant *ubiquitin*). Shown is the mean (±SD) from three biological replicates. Comparisons between’ groups were performed via ANOVA and Tukey’s Range Test. Letters represent statistical differences among all group means (α = 0.05).

### MoMIF1 is a secreted protein

In mammals, insect pests and parasitic protozoa, MIF proteins have been shown to be secreted molecules that enable parasitism and suppress immune responses and cell death (Augustijn et al., 2007; Vermeire et al., 2008; Naessens et al., 2015). To determine whether *Mo*MIF1is a secreted protein, we collected intercellular washing fluid (IWF) from WT and *MoMIF* OE mutant-infected leaves at 2 dpi for proteomics analysis. The IWFs were digested in solution with trypsin/lys-C enzymes and the resulting peptides were analyzed by ultra-high-performance liquid chromatography high-resolution tandem mass spectrometry (UHPLC–HR-MS/MS). Four unique peptides were identified with high confidence and confirmed as *Mo*MIF1 protein just in the IWF belonging to Δ*Mo*MIF1-OE_1-infected leaves (Fig. S6). On the contrary, no *Mo*MIF1 peptide was detected in the IWFs of WT-infected leaves. The latter is consistent with the low amount of *Mo*MIF1 at the 2 dpi time point as anticipated from its expression profile (see Fig. 2a). Importantly, we could not detect any peptides related to broken mycelial remnants (tubulin or actin proteins), suggesting that the *Mo*MIF1 was secreted by the OE mutant during the interaction with the plant. Furthermore, no *Mo*MIF1 peptide was detected in the sterile culture filtrates of either the OE mutants or the WT after 5 dpi, suggesting that the *Mo*MIF1 protein is secreted only when *Mo* is in contact with its host.

### MoMIF1 has cell death inhibitory activity

MIF proteins inhibit apoptosis and autophagy-dependent cell death in mammals, insects, and plants (Naessens et al., 2015; Balogh et al., 2018; Zhao et al., 2019; Zhao et al., 2021). We therefore performed Agrobacterium-mediated transient expression assays using leaves of *N. benthamiana* (Nb) to analyze whether *Mo*MIF1 has cell death inhibitory activity. Bcl-2-associated protein X (BAX) activates apoptotic cell death in mammals (Boise et al., 1995), which is inhibited by mammalian MIF (Baumann et al., 2003). Mammalian BAX also triggers hypersensitive response-like cell death in plant tissues upon transient expression of its gene (Lacomme and Santa Cruz, 1999; Hückelhoven et al., 2003). To analyze whether *Mo*MIF1 can inhibit BAX-induced cell death in plants, we performed *Agrobacterium tumefaciens*-mediated transient co-expression experiments in *Nb* leaves. Expression of murine (*Mm*) *BAX* (with *GFP* as control) gave rise to the expected cell death phenotype of the *Agrobacterium*-infiltrated leaf area (Fig. S7a-c; top-left half of the leaf), whereas expression of *MoMIF1* (with the *GFP* control) did not affect tissue integrity (Fig. S7a-c; bottom-left half of the leaf). Remarkably, when *MmBAX* and *MoMIF1* were simultaneously expressed in the same leaf area, the *Mm*BAX-induced cell death area was clearly and significantly smaller compared to *MmBAX* alone (Fig. S7a-c; right half of the leaf). These data together suggest that the fungus secretes *Mo*MIF1 to inhibit host cell death during the biotrophic phase of host colonization.

### MoMIF1 regulates the production of tenuazonic acid

Regulation and fine-tuning of developmental processes is characteristic of chemokines (Bromley et al., 2008). Studies with animal cells during developmental and pathological processes have revealed links between MIF proteins and the regulation of primary and secondary metabolic processes (Mitchell et al., 2002; Serre-Beinier et al., 2010). We therefore tested whether *MoMIF1* WT and KO mutants differ in secondary metabolite production. Chemical analysis by means of HPLC and HPLC-MS revealed that the KO mutants produced increased amounts of tenuazonic acid (TeA), independent of the nature of the culture medium, whereas the WT produces only traces thereof (Fig. 6a-b). TeA is a well-known non-host selective mycotoxin that inhibits protein biosynthesis at the ribosome by suppressing the release of new proteins. It is particularly important for the establishment of infection by many necrotrophic fungi, such as those from the genera *Alternaria* and *Phoma*, that secrete (TeA), thereby causing ROS production necroses and cell death in infected leaves (Yun et al., 2015; Shi et al., 2021). Overall, these results are consistent with the notion that *Mo*MIF1 is a negative regulator of TeA production, suggesting a scenario in which transient *Mo*MIF1 expression at early infection stages inhibits the mycotoxin production during the biotrophic phase, while subsequent downregulation of *Mo*MIF1 de-represses its production in the necrotrophic phase.

**Fig. 6.**
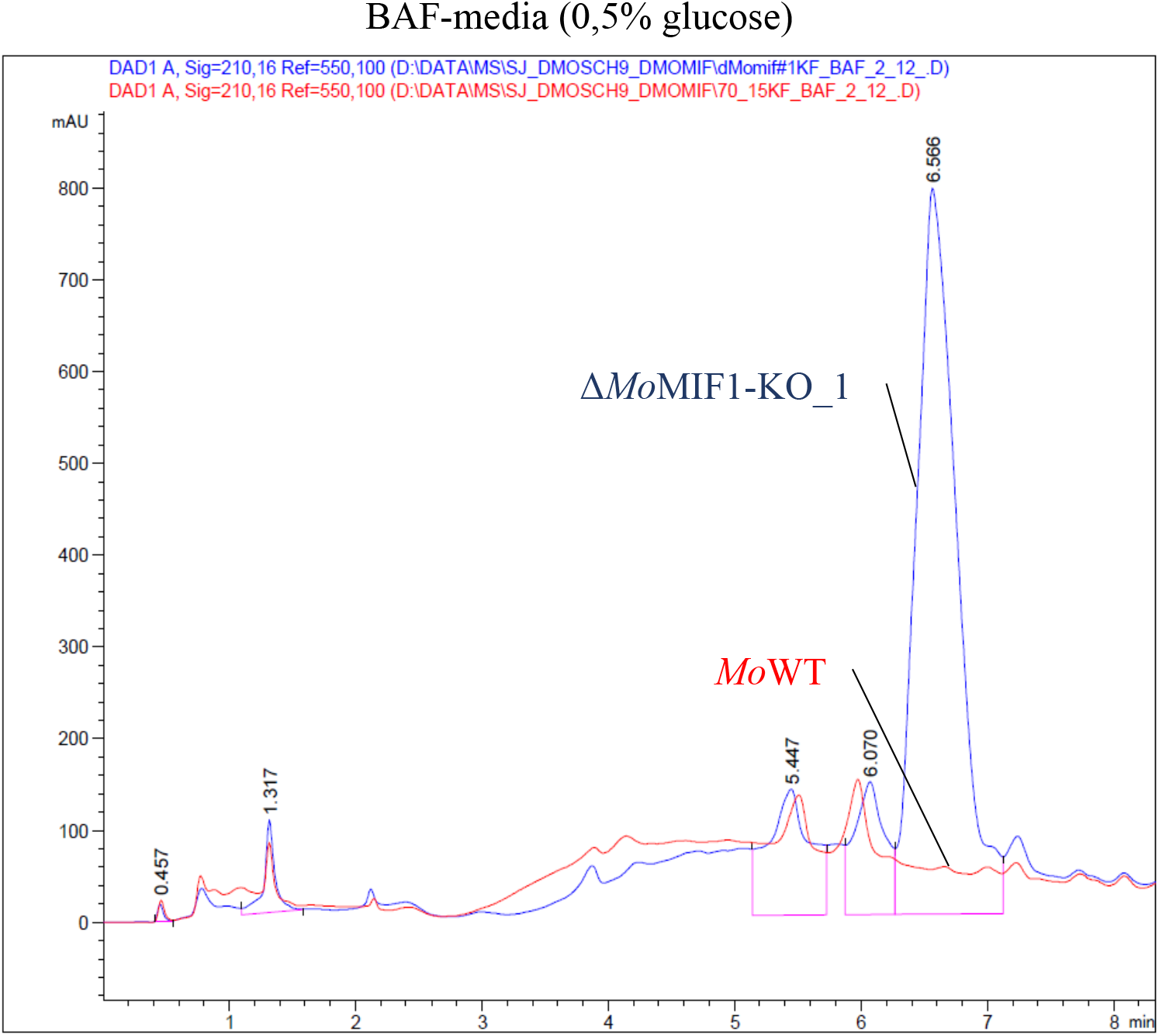
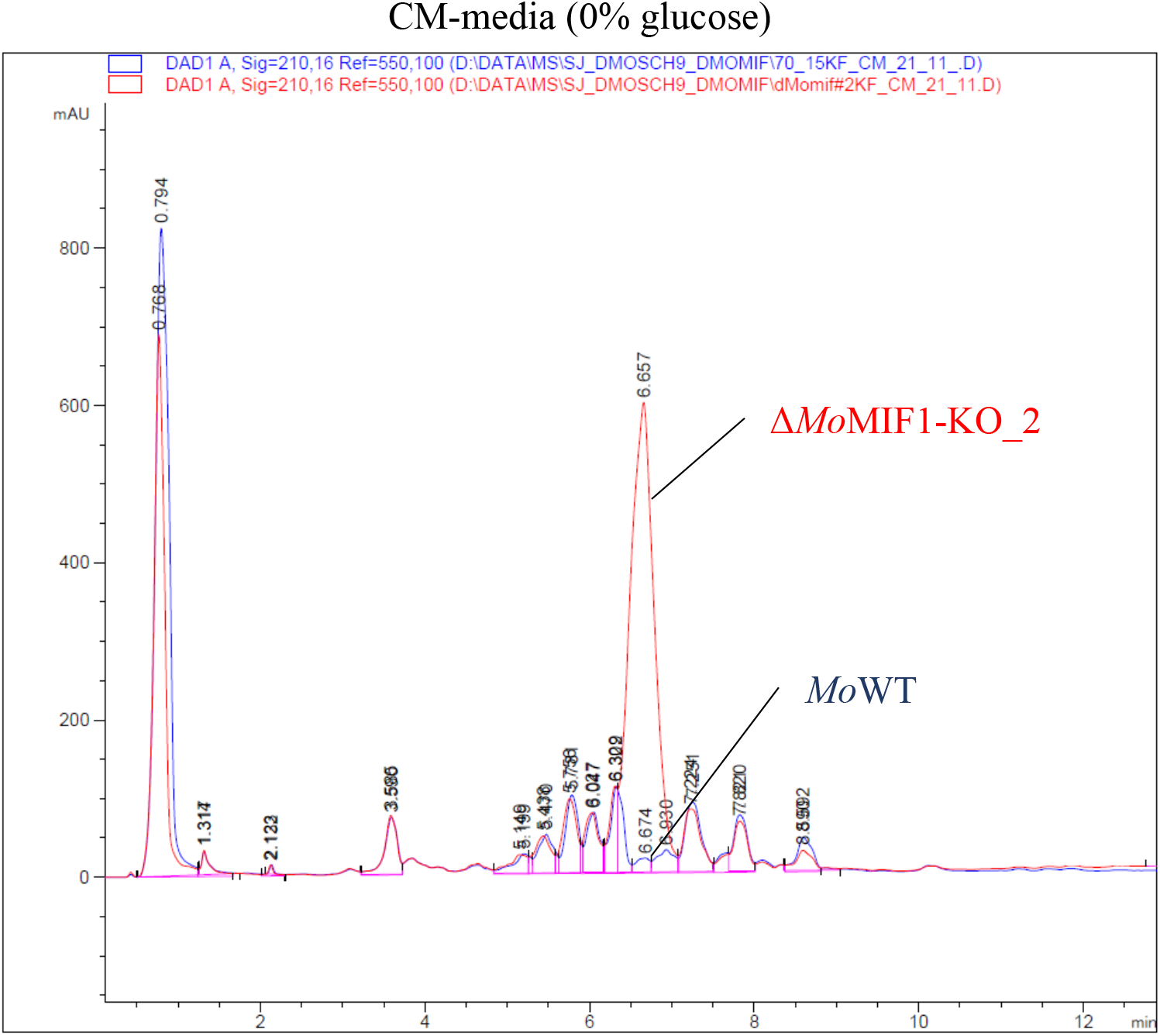
Quantification of tenuazonic acid (TeA) in *Mo*WT and MIF1 mutants. HPLC-MS analysis of metabolites extracted from WT and the KO mutant Δ*Mo*MIF1-KO_1 and Δ*Mo*MIF1-KO_2 in BAF medium plus 0.5 % Glucose (a) or CM (b). The indicated peaks mark the TeA area. Each strain was submerged-cultured for 5 days at 25°C in axenic cultures. The experiment was repeated three times with similar results.

## Discussion

Based on a survey of MIF homologues in the kingdom Fungi (EnsemblFungi database; Howe et al., 2020), we discovered that members of the monophyletic group Ascomycota carry atypical MIF proteins, which are unique to this phylum (Table 1; Fig S1). Compared to the closer phyla Mucoromycota, Zoopagomycota, Glomeromycota and Basidiomycota, which harbor typical MIF proteins, Ascomycota carry larger MIFs, in which the tautomerase domain is flanked by long peptide “arms” (Fig. 1a). Similarly, the marine dinoflagellate *Lingulodinium polyedrum* (Lp) from the phylum Myzozoa (infrakingdom Alveolata) also produces an atypical MIF protein (Jaouannet et al., 2020). LpMIF (246 aa) is a transmembrane-anchored MIF protein with an extracytoplasmic MIF domain associated with vesicle bodies, cell wall and extracellular vesicles (Jaouannet et al., 2020). The presence of related sequences in such disparate phyla suggests that selective pressure has influenced the evolution of the MIF protein scaffold to meet different physiological needs. In Ascomycota, this hypothesis is specially backed by the presence of several LCR motifs with unknown function flanking the tautomerase/MIF domain (Fig. 1a). Recent experimental and computational evidence increasingly suggests that LCRs have adaptive and conserved roles in protein evolution (Ntountoumi et al., 2019; Mier et al., 2020; Dickson and Golding, 2022). In particular, they may contribute to protein conformational plasticity therefore broadening the range of interacting molecules (Ntountoumi et al., 2019).

We discovered that a fungal MIF is a highly regulated gene whose activity strictly correlates with the two parasitic life stages of the hemibiotrophic fungus *Magnaporthe oryzae*, with transcripts accumulating in the early germination/penetration phase of biotrophic growth and dropping off sharply as it enters necrotrophic growth on the host plant (Fig. 2a). Remarkably, there is practically no gene expression detected in the necrotrophic growth phase, neither in the infected leaf nor in the root tissue. As we found that the *Mo*MIF has cell death inhibitory activity (Fig. S7), it is plausible that high gene expression in this host-cell death associated phase (Sesma and Osbourn, 2004; Tucker et al., 2010) would be detrimental to fungal development and virulence. Thus, our data together suggest that accumulation of *Mo*MIF1 is required at stages where the pathogenic fungus is confronted with the very early host immune response, i.e. during the attachment and colonization of the host surface and during the biotrophy-associated growth within the first leaf cell, where it functions as cell death inhibitor and negative regulator of the mycotoxin TeA, thus keeping host cells alive (Fig. 6a,b). The detection of *Mo*MIF1 in intercellular washing fluids at an early stage of plant infection (Fig. S6) further supports this scenario.

*In-planta* experiments with fungal mutants carrying aberrant *MIF1* genes further highlighted the importance of a strict and finetuned regulation of *MoMIF1* expression for the development of the typical blast symptoms as both KO and OE mutants are strongly attenuated in virulence (Fig. 2; Fig 3a-c; Fig. S4). Excitingly, bypassing the appressorium-mediated entry and biotrophic epidermal cell invasion by injecting *Mo* conidia directly into the mesophyll apoplastic space, rendered KO mutants highly aggressive, while higher amounts of *Mo*MIF1 in *MoMIF1* OE mutants reduced the spread of infection (Fig. 4; Fig. S5; Fig. 5a,b), unambiguously demonstrating that *Mo*MIF activity is detrimental to fungal virulence in the necrotrophic phase.

*Mo* secretes effector proteins and structurally distinct secondary metabolites into the host to suppress plant immunity, manipulate the biological activities and finally kill the plant cell (Thines et al., 2004; Collemare et al., 2008; Wilson and Talbot, 2009). We discovered that *Mo*MIF1 is a negative regulator of tenuazonic acid, a host cell death-inducing mycotoxin secreted by necrotrophic fungi such as those from the genus Alternaria and Phoma (Yun et al., 2015; Shi et al., 2021), and which has also been described previously in the genus Magnaporthe (Aver’yanov et al., 2007). Interestingly, the latter authors reported the cell death-inducing activity of TeA in plants, but also that exogenous TeA reduces the fitness of the fungus. This shows that tight control of TeA biosynthesis is essential for fungal survival. Taken together, our findings are consistent with the view that in plant pathogenic organisms MIF proteins are important regulators of host cell death that serve to prolong parasitism and plant exploitation (Naessens et al., 2015; Balogh et al., 2018; Zhao et al., 2019; Zhao et al., 2021). *Mo* produces and secretes the atypical MIF1 protein to influence the infection life cycle of the pathogen through activities on both the fungus itself (metabolic reprogramming) and host plant cells (inhibition of cell death). We suggest that the protein contributes to the maintenance of biotrophy of the hemibiotrophic fungus, in analogy to what has been found for MIF activities in mammals and invertebrates.

## Acknowledgments

We thank Christina Birkenstock and Martina Claar for their excellent technical support. This work was carried out within the framework of the German/French cooperation program by the Deutsche Forschungsgemeinschaft (DFG) and the Agence Nationale de la Recherche (ANR) through project numbers Ko 1208/25-1 (KHK) and ANR-16-CE92-0014-01 (CC and HK), through the “Investments for the Future” LABEX SIGNALIFE: program reference ANR-11-LABX-0028-01 (CC and HK), as well as the DFG program RU5116 (KHK). PG and BS acknowledge financial support by DFG (INST 162/500-1 FUGG) and by the State of Hesse through LOEWE Center DRUID (Novel Drug Targets against Poverty-Related and Neglected Tropical Infectious Diseases). YZ was supported by German Academic Exchange Service (DAAD).

## Author contributions

MG, SJ and KHK wrote the manuscript; KHK, ET, MG, SJ, HK, CC and MG designed the study; MG, YZ, JI, VA, PG and WA prepared material for the experiments; KHK, CC, ET and BS provided methodology and instrumentation; MG, YZ, JI, VA, PG and WA conducted the experiments; MG, SJ, ET, CC, PG, HK and KHK analyzed all data and drafted the figures. All authors commented and reviewed the final manuscript.

## Supplement figure legends

**Fig. S1 |** Phylogenetic relationship and visualization of annotated features for various eukaryotic *MIFs*. (a) aa sequences of *Mo*MIF1 protein domain and *Hs*MIF domain homologs in other eukaryotic species were aligned using the MUSCLE algorithm and the evolutionary history was inferred by using the Maximum Likelihood method and Le’s and Gascuel’s model (Le and Gascuel, 2008). Domains were inferred via the online-tool SMART with PFAM (Letunic et al., 2021). The maximum likelihood tree was generated using the MEGA software (MEGA X version 10.0.5, Kumar et al., 2018). Numbers in the tree nodes indicate percent confidence values based on 1000 bootstrap replicates. The tree is drawn to scale, with branch lengths measured in the number of substitutions per site (next to the branches). *Mo* is marked with a red square.

**Fig. S2 |** Generation of fungal strains used in this study. (a) Schematic representation of the cloning strategy and the genomic DNA of the WT strain and the mutant strain Δ*Mo*MIF1 and southern blot analysis with specific probes. The pink marked primer pairs marked the position of the hybridization of the probes. Genomic DNA of *Mo* strain 70-15 and the *MoMIF1* (MGG_05693) mutants were isolated and restricted with *BstBI* restriction enzymes. The probes which we used for hybridization with the genomic DNA of the wild□type strain and the corresponding mutant strains were always identical. (b) *MoMIF1* mRNA accumulation was quantified in WT, two KO mutants, two OE mutants, and the compensation mutant at 5 dpi in fungal mycelia grown in sterile condition via RT-qPCR. *MoActin* was used as the normalization gene. Shown is the average of three biological repetitions. Bars represent the standard deviation. Comparisons between groups were performed via ANOVA and Tukey range test for multiple comparisons. Letters represent statistical differences among all group means (α < 0.05).

**Fig. S3 |** Appressoria cytorrhysis assay and phenotypical analysis of axenic cultures of *Mo* mutants. (a) 5 × 10^3^ ml^-1^ conidia in distilled water were inoculated onto hydrophobic coverslips and appressoria formation was induced via 1,16-Hexadecanediol (500 ng/ml). 16 hpi distilled water was replaced with different glycerin solution and collapsed appressoria were counted after 5 minutes. Bars represent the standard deviation (±SD) of three biological repetitions (n=100 appressoria). (b) Osmotic stress test on different media. Agar blocks of approximately 0.8 cm diameter from 11-day-old *Mo* cultures grown on CM were cut out to be transferred onto CM or MM agar plates with different stress-inducing compounds (sorbitol, KCl, NaCl). The axenic cultures (n=3) were grown for 10 days at 26°C and diameters of the mycelia measured in millimeters [mm]. In this experiment *Mo*WT and two independent KO mutants (Δ*Mo*MIF1-KO_1, Δ*Mo*MIF1-KO_2) were used. Comparisons between groups were performed via ANOVA and Tukey range test for multiple comparisons. Letters represent statistical differences among all group means (α < 0.05). Glc: glucose, M: molarity.

**Fig. S4 |** Outcome of spray inoculation of three-week-old *Oryza sativa*. *Oryza sativa* (cv. CO-39) seedlings with 5 x 10^4^ conidia ml^-1^ of WT and mutants. Graph displays number of necrotic lesions on the three most heavily infested rice leaves at 5 dpi. The experiment was conducted two times (n=12 plants) with similar results. Comparisons between groups were perform’d via ANOVA and Tukey’s Range Test. Letters represent statistical differences among all group means (α = 0.05).

**Fig. S5 |** Disease symptoms caused by *Mo*WT and MIF1 mutants on *Irachypodium distachyon* Bd21-3. Phenotypical analysis of infection phenotypes at 5 dpi on two-week-old seedling roots upon dip-inoculation with 1 ml suspension of 120 x 10^3^ conidia ml^-1^ in 0.002% Tween20. After *Mo* inoculation, plants were cultivated in artificial soil containing Vermiculite substrate (Deutsche Vermiculite GmbH, Sprockhövel, Germany) for 5 days in a growth chamber at 22°C/18°C (day/night cycle) with 60% relative humidity and a photoperiod of 16 h (240 μmol m^-2^ s^-1^ photon flux density).

**Fig. S6 |** Identification of *Mo*MIF1 in the intercellular washing fluid (IWF) of *Mo*-infected Bd21-3 leaves. The protein was identified using UHPLC-HR-MS/MS analysis of IWF of Δ*Mo*MIF1-OE_1-infected leaves at 2 dpi. Unique peptides were highlighted in green and identified with high confidence against the *Mo*MIF1 sequences.

**Fig. S7 |** Hypersensitive cell death detection in *Nb* leaves after agro-infiltration. *Nb* leaves (n=18) were simultaneously *Agrobacterium*-mediated transformed with vectors carrying cDNA sequence of *Mm*BAX + GFP (leaf top-left side), *Mo*MIF1 + GFP (leaf bottom-left side) and *Mm*BAX + *Mo*MIF1 (leaf right side). GFP was used as negative control for toxicity. Photos were taken 96 hpi under normal (a) and UV (b) light. Black arrows indicate location of agro-infiltration. (c) Size of the apoptotic area was assayed via ImageJ software at 96 hpi. Comparisons between groups were performed via Kruskal-Wallis Test with Dunn’s test with Bonferroni adjustment. Letters represent statistical differences among all group means (α = 0.05).

**Table S1 |** List of primers used in this study with relative target and function.

## STAR Methods

### Plant and fungal growth conditions and inoculation protocols

The fungal strain used in this study was *Magnaporthe oryzae* strain 70-15 strain (Fungal Genetics Stock Center, Kansas State University, Manhattan, USA). The strain was grown at 26°C on CM (pH 6.5, 2% agar, containing per liter 10 g glucose, 1 g yeast extract, 2 g peptone, 1 g casamino acids, 50 mL nitrate salt solution [120 g NaNO_3_, 10.4 g KCl, 30.4 g KH_2_PO_4_, 10.4 g MgSO_4_ x 7 H_2_O per liter] and 1 mL of a trace element solution [22 g ZnSO_4_ x 7 H_2_O, 11 g H_3_BO_3_, 5 g MnCl_2_ x 4 H_2_O, 5 g FeSO_4_ x 7 H_2_O, 1.7 g CoCl_2_ x 6 H_2_O, 1.6 g CuSO_4_ x 5 H_2_O, 1.5 g Na_2_MoO_4_ x 2 H_2_O and 50 g Na_2_EDTA, pH 6.5 adjusted by 1 M KOH per liter]. Minimal medium (MM; pH 6.5) contains 1 g glucose, 0.25 mL of a 0.01% biotin solution, 50 mL nitrate salt solution, 1 mL of a trace element solution, and 1 mL of a 1% thiamine dichloride solution per liter. Germination of conidia and appressoria development were assessed by incubating a suspension of 5 x 10^3^ conidia ml^−1^ in double distilled water on plastic coverslips (Sigma-Aldrich, Saint Louis, USA) in a damp chamber at room temperature fIsing an inverted microscopy. *Brachypodium distachyon* cv. Bd21-3 (Vogel and Hill, 2008) was cultivated in soil (Fruhstorfer Erde Typ T, Vechta, Germany) in a growth chamber at 22°C/18°C (day/night cycle) with 60% relative humidity and a photoperiod of 240 μmol m^-2^ s^-1^ photon flux density. Dwarf indica rice cultivar CO-39 was cultivated using a daily cycle of 16 h/8 h light/dark (28°C, 80% relative humidity). Two methods were used to assess disease progression of *Mo* strains in plant leaves: *i*. Leaf-spray inoculation of three-week-old Bd21-3 plants with a suspension of 120 x 10^3^ ml^-1^ conidia in 0.002% v/v Tween20 until water run-off; *ii*. injection of 100 μl of the above suspension into the apoplastic space of second youngest intact leaves of three-week-old Bd21-3 using a syringe. Tenuazonic acid (TeA, copper salt, Cayman Chemical Company, Michigan, USA) was used at a concentration of 5 mM in double distilled H_2_O (TeA first resuspension in 0.5% v/v DMSO). Control seedlings were mock-inoculated with 0.002% v/v Tween20 water in all setups. Disease progression and analysis of the necrotic spots were assayed at 5 dpi via ImageJ software. For rice infection, conidial suspensions were adjusted to 5 x 10^4^ conidia ml^−1^ in H_2_O containing 0.2% gelatin. Five 21-day-ol rice plants were spray inoculated each with 5 mL of conidial suspension and incubated in plastic bags in a test chamber (Versatile Environmental Test Chamber MLR-350H; Sanyo Electric Co., Illinois, USA). For root inoculation, Bd21-3 seeds were surface sterilized with a solution of 3% v/v NaClO for 15 min, followed by three times 5 min washes in sterile water, before being placed on half-strength MS medium in 16 h light (160 μmol□m^-2^ s^-1^)/8 h dark cycle at 22°C/18°C. Roots of one-week-old seedlings were dip-inoculated in 1 ml of conidial suspension (120 x 10^3^ ml^-1^ in 0.002% v/v Tween20) for 3 h and later transplanted in vermiculite (Deutsche Vermiculite GmbH, Sprockhövel, Germany). Control roots were mock inoculated with 1 ml 0.002% v/v Tween20. At 5 dpi, fungal DNA was extracted from roots and leaves using a DNA extraction kit (Qiagen, Hilden, Germany), and quantified by RT-qPCR using fungal *actin* (*MoActin*) or *glyceraldehyde-3-phosphate dehydrogenase* (*MoGPD*) and normalized to *Bd ubiquitin* (*BdUbiquitin*) (Table. S1) as described in Zanini et al. (2021).

### Statistics

ANOVA test with Pairwise Comparisons with Bonferroni adjustment, ANOVA with posthoc Tukey HSD test or Kruskal Wallis with Dunn’s test with Bonferroni adjustment were selected after analysis of the sample distribution in the different groups.

### DNA manipulations and construction of fungal transformation vectors

The DNA of *Mo* 70–15 was isolated from mycelia of 3-day-old liquid cultures (grown in CM at 26°C and 100 rpm) using the DNeasy^®^ plant mini Kit (Qiagen GmbH, Hilden, Germany), following the manufacturer’s instructions for purification of DNA from plants and filamentous fungi. The DNA manipulation followed standard procedures. *Escherichia coli* XL1-BLUE strain (Stratagene) or NEB^®^ 10-β Competent *Escherichia coli* strains (High Efficiency) were used for routine bacterial transformations and the construction of plasmids. Transformation of *Mo* was conducted using *Agrobacterium tumefaciens*-mediated transformation (ATMT). The detailed procedures followed those described in Odenbach et al. (2007).

To generate the MIF1 loss-of-function mutants Δ*Mo*MIF1-KO_1 and Δ*Mo*MIF1-KO_2, the coding gene sequence of *MoMIF1* (MGG_05693) was completely replaced by the HPT gene from *pSJ-basic* (IBWF-73, Bohnert et al., 2019), using the Gibson Assembly^®^ cloning method. The flanking regions (1000 bp) of *MoMIF* were amplified by PCR from genomic DNA with primers SJ-1249/SJ-1250 and SJ-1253/SJ-1254. The *HPT* gene was amplified from *pSJ+basic* using the primer SJ-1251/SJ-1252. The backbone vector was a *PstI/BglII-restricted pSJ-basic*. The 7,917 bp plasmid *pSJ-MIF-HPT* was used to transform WT via ATMT, resulting in the mutant strain Δ*Mo*MIF1-KO_1 and Δ*Mo*MIF1-KO_2.

For the fluorescence marked strain Δ*Mo*MIF1+GFP, we used a two-step cloning strategy. We initially generated the plasmid *pSJ+flank1+EF1+GFP+BAR+flank2*. Therefore, the plasmid *pSJ+GFP*(*BAR*) (Bohnert et al., 2019) was used as template to amplify fragment *EF1+GFP+BAR* using the primer pair SJ-1628/SJ-1629. Fragment 1 is generated in order to obtain a strong expression of the *eGFP*-gene, so the *EF1α*-promoter of *Mo* (gene MGG_03641) was cloned N-terminal of the *eGFP* gene, which was originally amplified from *pEGFP-1* (BD Biosciences). At the C-terminal end of the *eGFP* sequence, the *NOS*-terminator was inserted, directly followed by the *BAR* sequence (Bohnert et al., 2019). The flanking regions (500 bp) of *MoMIF1* were amplified by PCR from genomic DNA with primers SJ-1626/SJ-1627 and SJ-1630/SJ-1631. The backbone vector was a *PstI/BglII*-restricted *pSJ-basic*. The resulting 8,562 bp plasmid *pSJ+flank1+EF1+GFP+BAR+flank2* was then used as backbone to insert the MIF coding sequence in the *Nco*I restriction site between *EF1* and *GFP*. The MIF fragment was amplified by PCR from genomic DNA with primers SJ-1632/SJ-1633. The 9,958 bp plasmid *pSJ+flank1+EF1+MIF+GFP+BAR+flank2* was used to transform Δ*Mo*MIF1-KO via ATMT, resulting in the mutant strain Δ*Mo*MIF1+GFP.

For the overexpression strain Δ*Mo*MIF1-OE_1, the *Nco*I/*Bsr*GI-restricted plasmid *pSJ+flank1+EF1+GFP+BAR+flank2* was used as backbone for the cloning procedure. The *MoMIF1* coding sequence was amplified by PCR from genomic DNA with primers SJ-1634/SJ-1635. The 9,249 bp plasmid *pSJ+flank1+EF1+MIF+BAR+flank2* was used to transform Δ*Mo*MIF1-KO via ATMT, resulting in the mutant strain Δ*Mo*MIF1-OE_1.For the overexpression strain Δ*Mo*MIF1-OE_2, the *Hind*III*/Not*I-restricted plasmid *pSJ+flank1+EF1+GFP+BAR+flank2* was used as backbone for the cloning procedure. To obtain a strong expression of the *MoMIF1*-gene, the promoter of *Mo* ribosomal protein 27 was amplified from plasmid pSM565 with the primers SJ-1636/SJ-1637. The *MoMIF1* coding sequence was amplified by PCR from genomic DNA with primers SJ-1638/SJ-1639. The 8,612 bp plasmid *pSJ+flank1+RP27+MIF+BAR+flank2* was used to transform Δ*Mo*MIF1-KO via ATMT, resulting in the mutant strain Δ*Mo*MIF1-OE_2. For the reintegration of the native *MoMIF1* gene to complement the strain Δ*Mo*MIF1-KO_1, the *PstI/BglII*-restricted *pSJ-basic* was used as backbone. The NOS+BAR fragment was amplified from *pSJ+GFP(BAR*) using the primer pair SJ-1642/SJ-1643. A 1,700 bp native promoter region with the following *MoMIF1* sequence was amplified from genomic DNA with primers SJ-1708/SJ-1709. Furthermore, the C-terminal flanking region (500 bp) of *MoMIF1* was amplified by PCR from genomic DNA with primers SJ-1726/SJ-1727. The resulting 9,279 bp plasmid *pSJ+natprom+MIF+BAR+flank2 was* used to transform Δ*Mo*MIF1-KO via ATMT, resulting in the complementation strain Δ*Mo*MIF1-comp.

### *Protein* organization *comparison and phylogenetic analysis*

Whole MIF and MIF domain amino acid sequences from selected species were used for the protein structure and phylogenetic analysis. Visualization and comparison of the different MIFs proteins were done via the online-tool CDD/SPARCLE (https://www.ncbi.nlm.nih.gov/Structure/cdd/wrpsb.cgi, Marchler-Bauer et al., 2017). Visualization and comparison of the different MIFs domains was performed via SMART with PFAM (http://smart.embl.de; Letunic et al., 2021) Multiple sequence alignments were carried out using the MUSCLE algorithm (Edgar, 2004). The phylogenetic tree was built using the maximum likelihood statistical method based on the Le’s and Gascuel’s model (Le and Gascuel, 2008). Tree nodes accuracy was tested via the bootstrap method with 1000 bootstrap replicates. Phylogenetic and molecular evolutionary analyses were conducted using MEGA software (MEGA X version 10.0.5, Kumar et al., 2018). Corresponding coding and genomic sequences were obtained from the EnsemblFungi portal (https://fungi.ensembl.org/index.html, Howe et al., 2020).

### Production of sterile crude extracts from Mo cultures and intercellular washing fluid from Mo-infected Bd 21-3 leaves

To produce new axenic cultures three agar blocks of 10 mm diameter from 11-day-old *Mo* cultures were aseptically transferred as inoculum to 200 ml liquid CM in 500 ml glass flasks with one baffle. These cultures were grown at 26°C and 120 rpm. Intercellular washing fluid (IWF) was isolated from infected leaves as described (Pogorelko et al., 2011). Sterile solutions were obtained by filtration through disposable sterile filter (0.22 μm PES Membrane, Merck, Darmstadt, Germany). For HPLC-MS the solutions were dried with Na2SO4. After evaporation of the organic solvent *in vacuo* at 40°C to dryness, the residue was dissolved in methanol to give 10 mg/ml.

### HPLC/MS method for analysis of secondary fungal metabolites

Crude extracts were analyzed by means of HPLC (Agilent 1100 Series) equipped with a LiChrospher RP 18 (3×125 mm; 5 μm, Merck) and a diode array detector. In order to analyse the extracts, the temperature of the column was set to 40°C and a flow rate of 1 ml/min was used with an elution gradient composed of H_2_O and acetonitrile. The molecular weight of the peaks selected was determined using an HPLC-MS (Agilent 1260 Series LC and 6130 Series Quadrupole MS System). The mass spectra were recorded using atmospheric pressure chemical ionization in positive- and negative-ion mode. A Superspher RP 18 (125×2 mm; 4 μm, Merck) column was used at 40°C. For every run, 1 μl of a sample at a concentration of 1 mg/ml was injected. The elution was performed with a gradient of H2O and acetonitrile, and a flow rate of 0.45 ml/min.

### Protein purification and bottom-up proteomics analysis

The *Mo* supernatants were concentrated using tangential flow filtration (Ultrasette^™^ Lab Tangential Flow Device with Omega^™^ polyether sulfone membrane, Pall Deutschland Holding GmbH & Co. KG, Dreieich, Germany). The samples were prepared and digested before LC-MS/MS analysis as described in Ghezellou et al. (2021) and Sowa et al. (2022). In summary, samples were reduced and alkylated with 100 mM dithiothreitol (DTT) at 56 °C for 15 min and 200 mM iodoacetamide (IAA) at room temperature (dark place) for 30 min, respectively. The digestion was performed with a mass spec grade Trypsin/Lys-C mix (1:50 enzyme to proteins ratio; Promega GmbH, Mannheim, Germany) overnight.

The resulting peptides were separated by UltiMate 3000 RSLC UHPLC system (Thermo Fischer Scientific, Bremen, Germany) equipped with a Kinetix C18 column (Phenomenex, 2.6 μm, 100 Å, inner diameter 2.1 mm, 100 mm length) coupled with a Q Exactive HF-X Orbitrap (Thermo Fischer Scientific) mass spectrometer. Chromatographic analysis was performed at a flow rate of 250 μL min-1 with water/0.1% formic acid (mobile phase A) and acetonitrile/0.1% formic acid (mobile phase B). The gradient elution of 80 min was performed as follows: 2% B for 5 min, followed by 2-40% B over 60 min, 40-50% B over 5 min, 50-98% B over 5 min, and re-equilibration in 2% B. The mass spectrometer was operated in data-dependent acquisition (top-10 DDA) with the following parameters in full MS scans: a mass range of m/z 350-1800, mass resolution of 120,000 (@ *m/z* 200), AGC target of 3e6, injection time (IT) of 50 ms, and MS/MS scans: mass resolution of 30,000 (@ *m/z* 200), AGC target of 1e5, IT of 120 ms, isolation window *m/z* ±1.3, dynamic exclusion of 30 s and normalized collision energy (NCE) of 28. The raw files were processed using Proteome Discoverer version 2.4 (Thermo Scientific) with SEQUEST and MS Amanda search engines against the MIF databases.

### Vegetative growth assays

Stress tolerance of the mutant strains was conducted according to (Jacob et al., 2014). Agar blocks of approximately 0.8 cm diameter from 11-day-old *Mo* cultures grown on CM were cut out to be tested and transferred on CM or MM agar plates with different stress-inducing compounds (sorbitol, KCl, NaCl). The cultures were grown for 7 days at 26°C.

### Appressorium formation assays and conidia morphology

Appressorium development was assayed by monitoring germination of conidia on hydrophobic plastic slides (thickness 1, 76×51 mm, no. 653081; Greiner BioOne, Kremsmünster, Austria). For this purpose, conidia were harvested from 12-day-old *Mo* cultures grown on CM, filtered through two layers of miracloth to give a conidial suspension, which was adjusted to 5 x 10^4^ conidia mL^-1^ in ddH_2_O. 100 μl drops of this conidial suspension were placed on the plastic slides and incubated at room temperature. After 16 h the number of appressoria formed was counted. In addition, the same assay was carried out on glass slides upon chemical stimulation with 500 ng mL^-1^ 1,16-hexadecandiol (1,16-HDD, dissolved in MeOH). 1,16-HDD is a plant lipid or wax compound and an inducer of appressorium formation in *Mo* (Gilbert et al., 1996).

For the cytorrhysis assay, different concentrations of glycerol (1 M, 2 M, 3 M, 4 M) were added to the differentiated appressoria and after 5 min the number of collapsed appressoria was counted.

### Confocal laser scanning microscopy

Images were taken using a Leica TCS SP8 confocal laser scanning microscope (CLSM, Wetzlar, Germany) equipped with a 75-mW argon/krypton laser (Omnichrome, Chino, CA) and a water immersion objective (HCX APO L40×0.80 W U-V-l objective). Images were processed using the Leica LAS X software. GFP: green fluorescence λ_emission_: 508 nm; λ_excitation_: 489 nm).

### Transient expression in N. benthamiana

For transient expression analysis, *A. tumefaciens* GV3101 strains containing constructs with genes encoding either *Mo*MIF, mBAX, and GFP were grown in LB medium supplemented with 50 μg/mL rifampicin, 20 μg/mL gentamicin and 100 μg/mL spectinomycin until OD_600_ reached 1.0. Cells were pelleted, resuspended in infiltration buffer (10 mM MgCl_2_, 10 mM 2-[N-morpholino] ethanesulfonic acid [MES], pH 5.6, 200 μM acetosyringone) and adjusted to an OD_600_ of 0.6, then left for 3h at room temperature. The solutions were combined such that the final OD_600_ in the mixtures were 0.375 for *MoMIF-*, 0.125 for *GFP*-, and *0.05* for *mBAX*-containing agrobacteria. The abaxial side of leaves from 4-week-old *N. benthamiana* was infiltrated using a syringe without a needle. The leaves were scored for cell death 48 h after infiltration. For subcellular localization studies, *Mo*MIF1 was tagged at the N-terminus with eGFP (plasmid pK7WGF2) and transferred into Agrobacterium strain GV3101. The bacterium was grown as described above and infiltrated into *N. benthamiana* leaves at on OD_600_ of 0.5. Leaf patches were analyzed by CLSM 48 h after infiltration on an inverted Zeiss LSM880 microscope (Carl Zeiss France SAS, Marly-le-Roi, France), equipped with Argon ion and HeNe lasers as excitation sources. Samples were excited at 488 nm and images were acquired with a C-Apochromat 63x/1.20 W Corr M27 objective and processed using the Zeiss ZEN 2 software package.

