## supplement table 1 for "A key regulator with a MIF-like domain orchestrates cellular differentiation and virulence in the fungal pathogen *Magnaporthe oryzae*"

Fig S1a

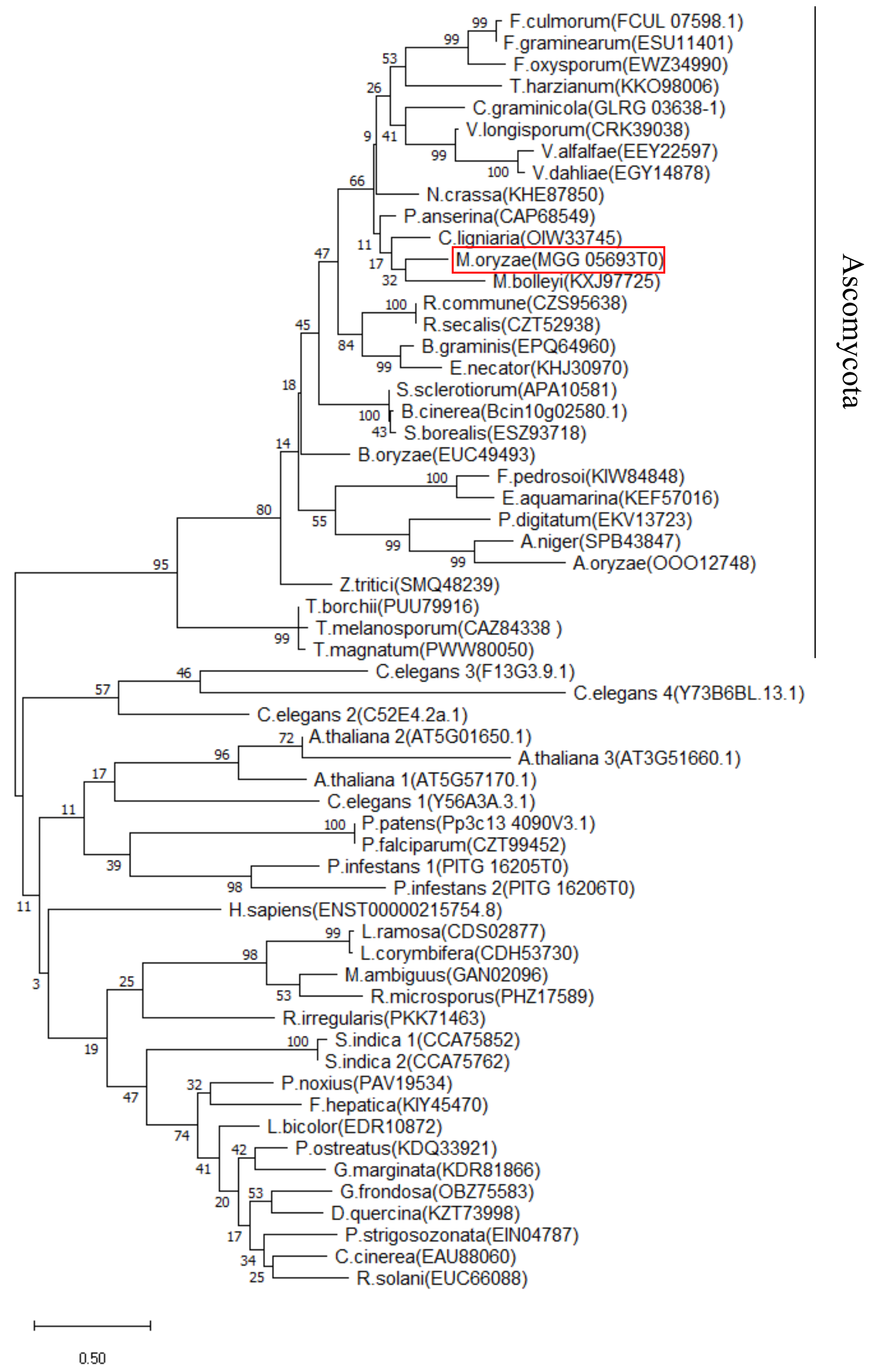
F.culmorum(FCUL 07598.1)
F.graminearum(ESU11401)
F.oxysporum(EWZ34990)
T.harzianum(KKO98006)
C.graminicola(GLRG 03638-1)
V.longisporum(CRK39038)
V.alfalfae(EEY22597)
V.dahliae(EGY14878)
N.crassa(KHE87850)
P.anserina(CAP68549)
C.ligniaria(OIW33745)
M.oryzae(MGG 05693T0)
M.bolleyi(KXJ97725)
R.commune(CZS95638)
R.secalis(CZT52938)
B.graminis(EPQ64960)
E.necator(KHJ30970)
S.sclerotiorum(APA10581)
B.cinerea(Bcin10g02580.1)
S.borealis(ESZ93718)
B.oryzae(EUC49493)
F.pedrosoi(KIW84848)
E.aquamarina(KEF57016)
P.digitatum(EKV13723)
A.niger(SPB43847)
A.oryzae(OOO12748)
Z.tritici(SMQ48239)
T.borchii(PUU79916)
T.melanosporum(CAZ84338 )
T.magnatum(PWW80050)
Ascomycota
C.elegans 3(F13G3.9.1)
C.elegans 4(Y73B6BL.13.1)
C.elegans 2(C52E4.2a.1)
A.thaliana 2(AT5G01650.1)
A.thaliana 3(AT3G51660.1)
A.thaliana 1(AT5G57170.1)
C.elegans 1(Y56A3A.3.1)
P.patens(Pp3c13 4090V3.1)
P.falciparum(CZT99452)
P.infestans 1(PITG 16205T0)
P.infestans 2(PITG 16206T0)
H.sapiens(ENST00000215754.8)
L.ramosa(CDS02877)
L.corymbifera(CDH53730)
M.ambiguus(GAN02096)
R.microsporus(PHZ17589)
R.irregularis(PKK71463)
S.indica 1(CCA75852)
S.indica 2(CCA75762)
P.noxius(PAV19534)
F.hepatica(KIY45470)
L.bicolor(EDR10872)
P.ostreatus(KDQ33921)
G.marginata(KDR81866)
G.frondosa(OBZ75583)
D.quercina(KZT73998)
P.strigosozonata(EIN04787)
C.cinerea(EAU88060)
R.solani(EUC66088)
0.50

Fig S2a

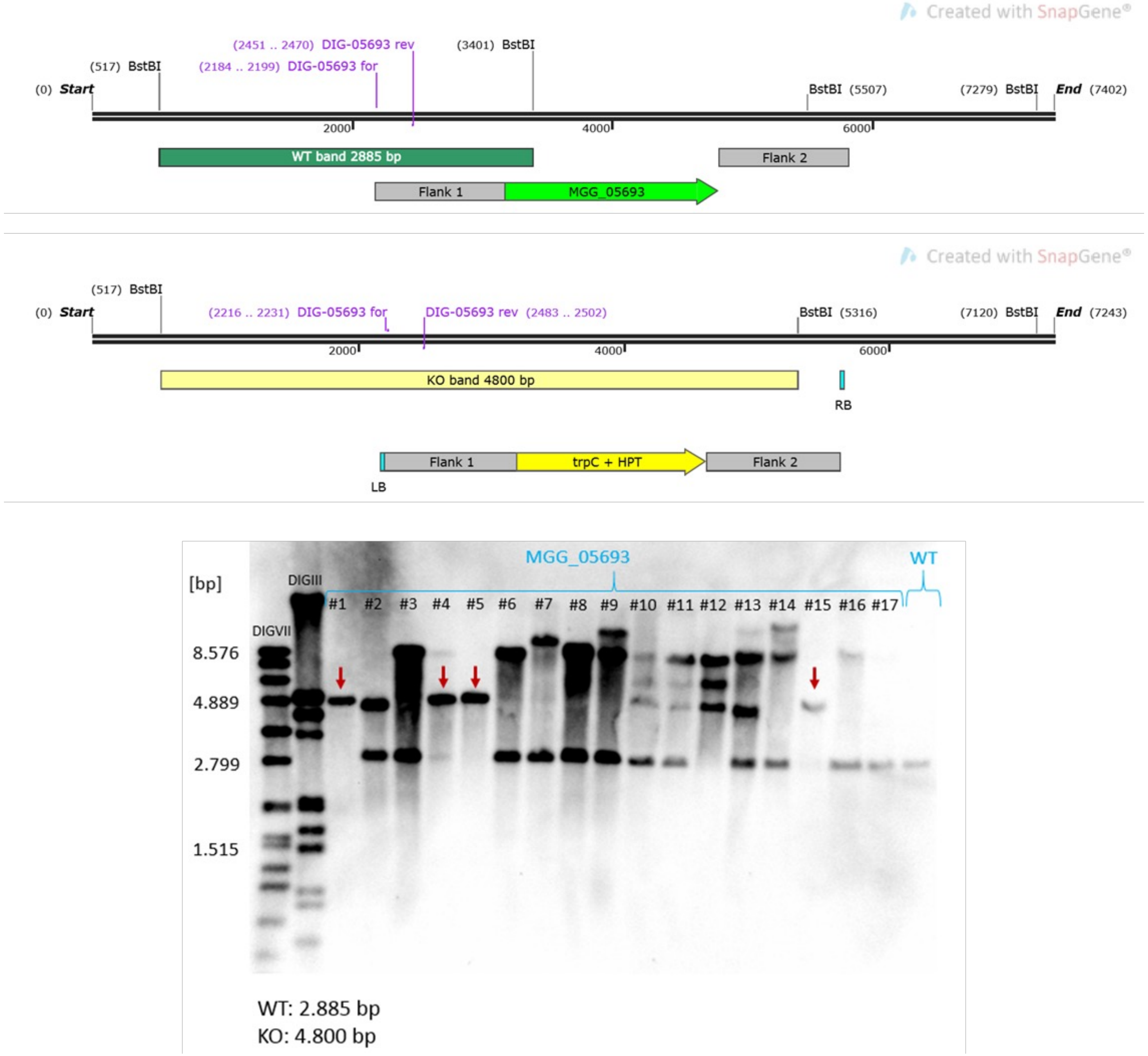
Created with SnapGene®
(2451 .. 2470) DIG-05693 rev
(3401) BstBI
(517) BstBI
(2184 .. 2199) DIG-05693 for
(0) Start
BstBI (5507)
(7279) BstBI
End (7402)
2000
4000
6000
WT band 2885 bp
Flank 2
Flank 1
MGG_05693
Created with SnapGene®
(517) BstBI
(0) Start
(2216 .. 2231) DIG-05693 for
DIG-05693 rev (2483 .. 2502)
BstBI (5316)
(7120) BstBI
End (7243)
2000
4000
6000
KO band 4800 bp
RB
Flank 1
trpC + HPT
Flank 2
LB
MGG_05693
WT
[bp]
DIGIII
DIGVII
#1 #2 #3 #4 #5 #6 #7 #8 #9 #10 #11 #12 #13 #14 #15 #16 #17
8.576
4.889
2.799
1.515
WT: 2.885 bp
KO: 4.800 bp

Fig S2b

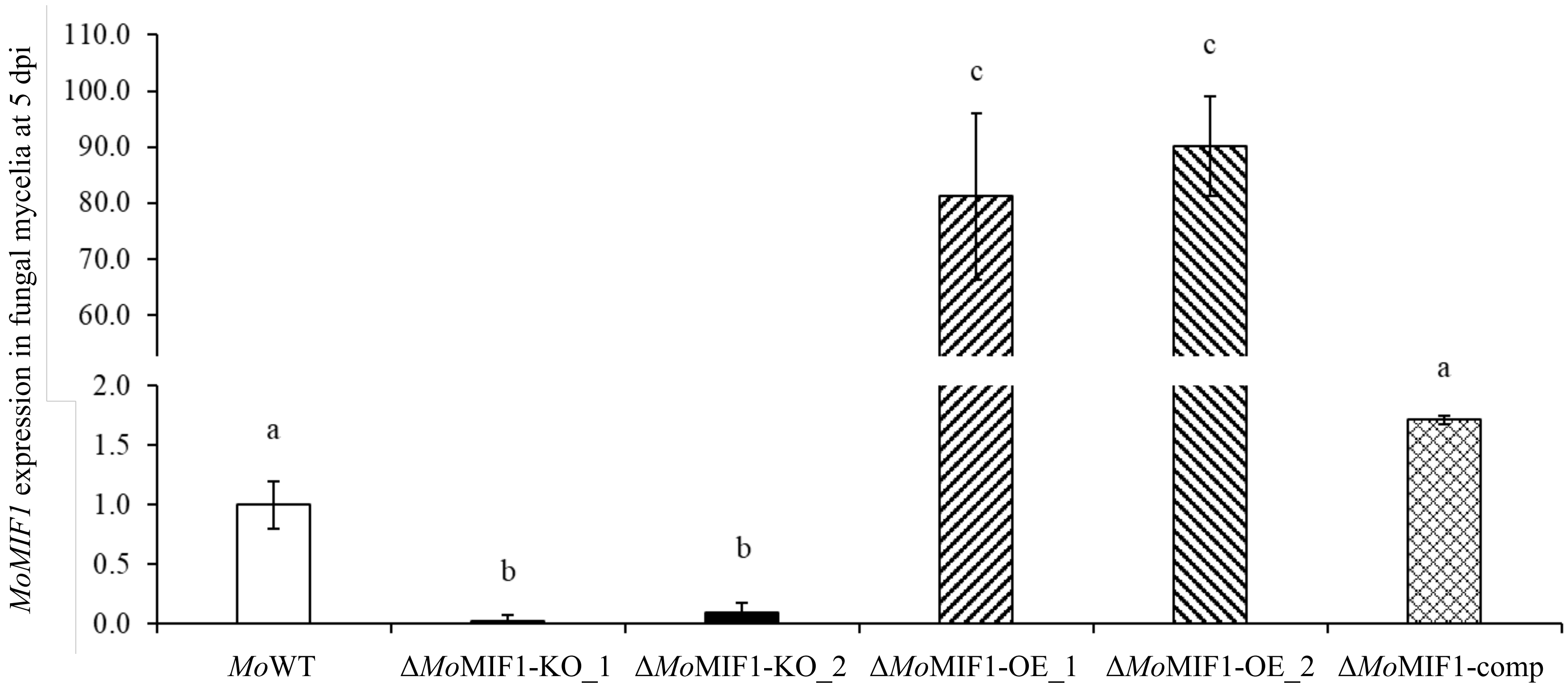

MoMIF1 expression in fungal mycelia at 5 dpi
110.0
100.0
90.0
80.0
70.0
60.0
2.0
1.5
1.0
0.5
0.0
a
b
b
c
c
a
MoWT
ΔMoMIF1-KO_1
ΔMoMIF1-KO_2
ΔMoMIF1-OE_1
ΔMoMIF1-OE_2
ΔMoMIF1-comp

Fig S3a

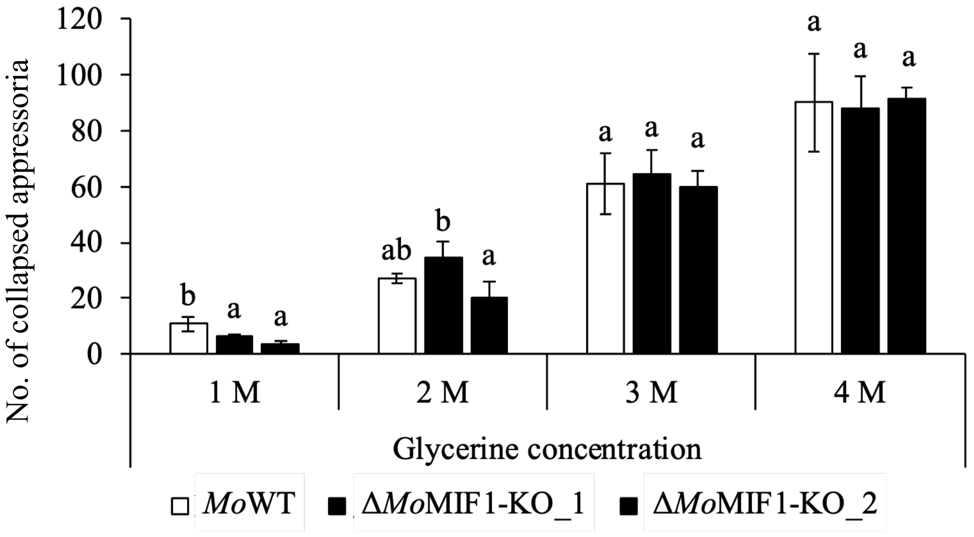

120
100
80
60
40
20
0
No. of collapsed appressoria
b
a
a
ab
b
a
a
a
a
a
a
a
1 M
2 M
3 M
4 M
Glycerine concentration
*Mo*WT
Δ*Mo*MIF1-KO_1
Δ*Mo*MIF1-KO_2

Fig S3b

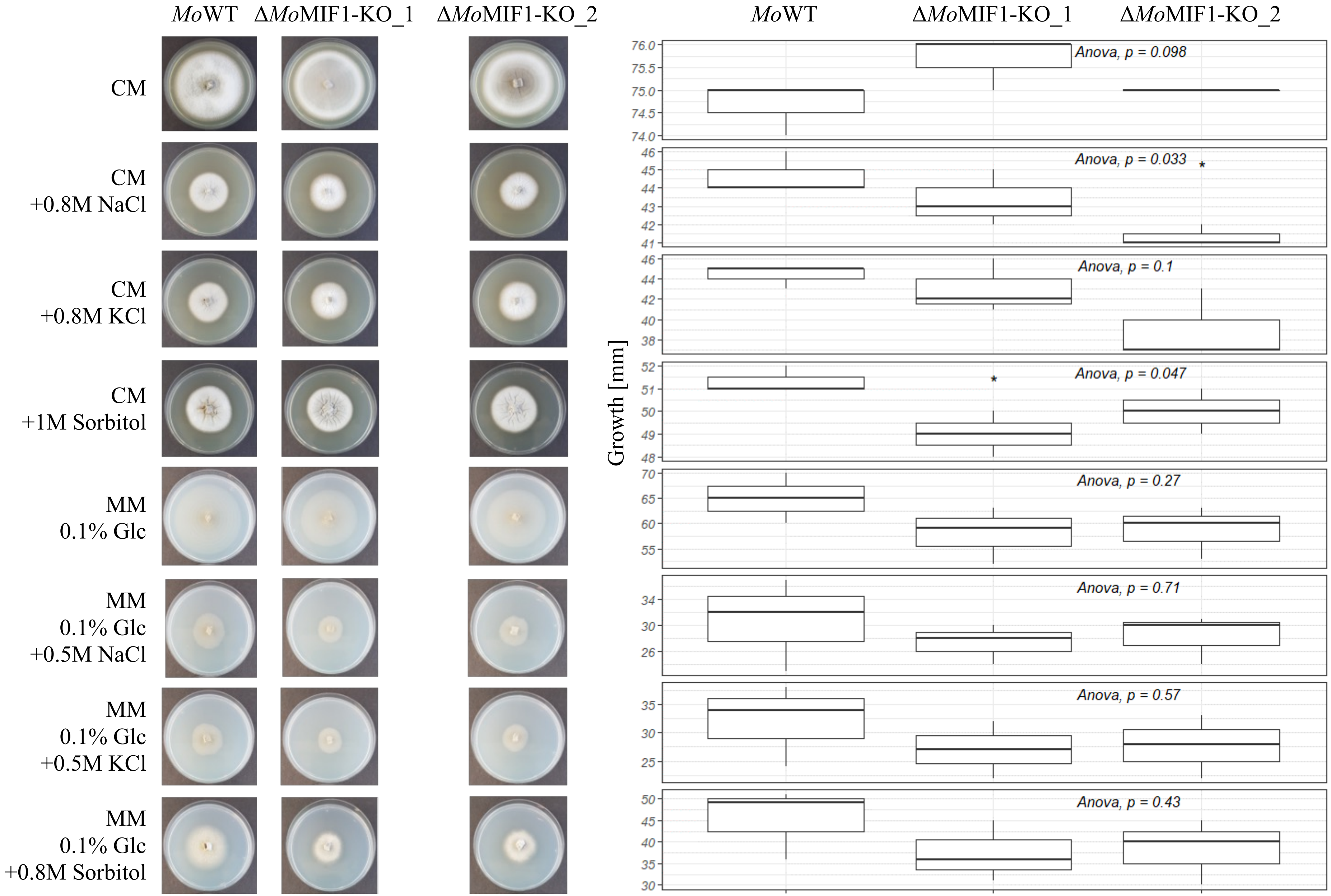
MoWT
ΔMoMIF1-KO_1
ΔMoMIF1-KO_2
CM
CM
+0.8M NaCl
CM
+0.8M KCl
CM
+1M Sorbitol
MM
0.1% Glc
MM
0.1% Glc
+0.5M NaCl
MM
0.1% Glc
+0.5M KCl
MM
0.1% Glc
+0.8M Sorbitol
Growth [mm]
Anova, p = 0.098
Anova, p = 0.033
Anova, p = 0.1
Anova, p = 0.047
Anova, p = 0.27
Anova, p = 0.71
Anova, p = 0.57
Anova, p = 0.43

Fig S4

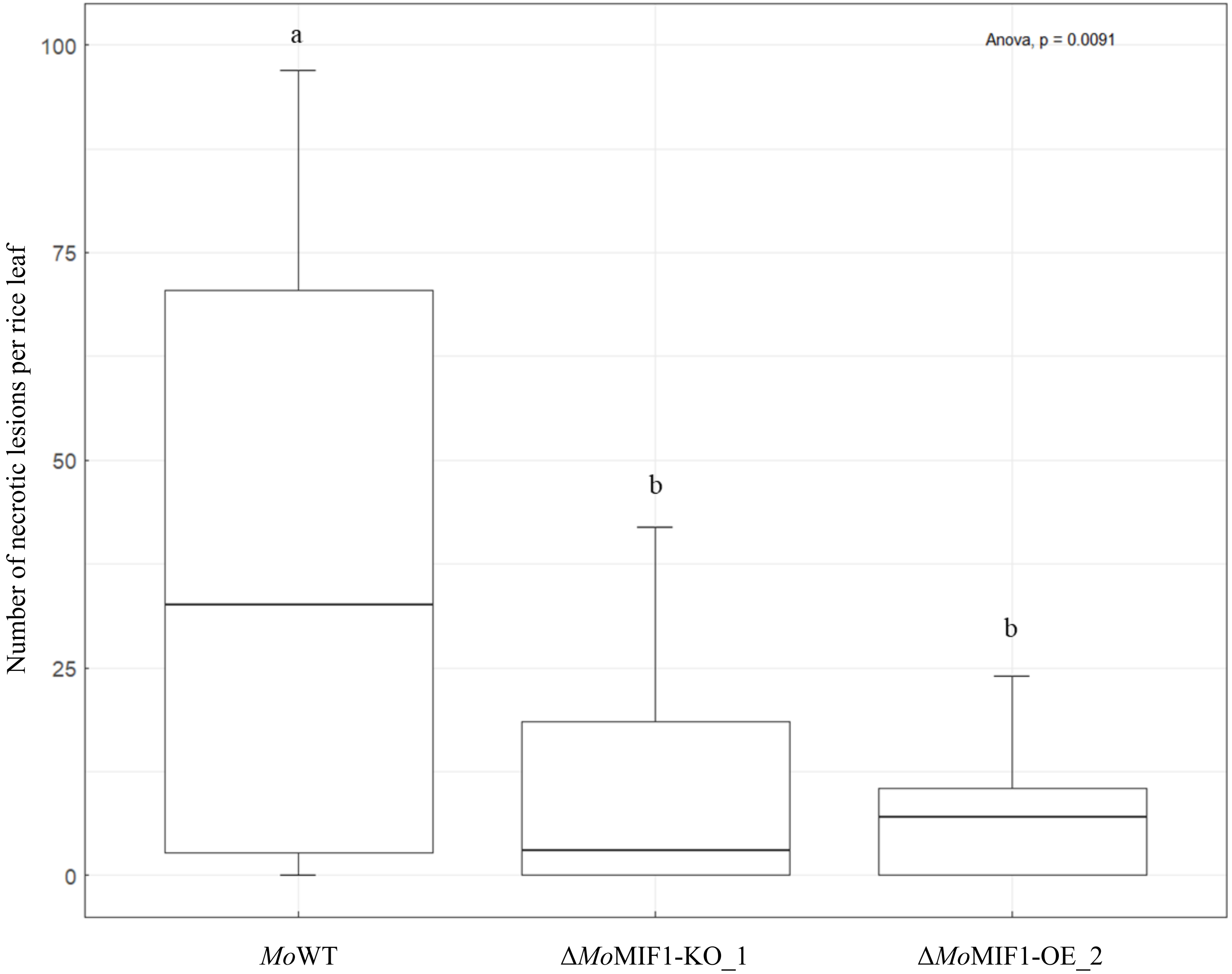

Number of necrotic lesions per rice leaf
100
75
50
25
0
a
b
b
Anova, p = 0.0091
*Mo*WT
Δ*Mo*MIF1-KO_1
Δ*Mo*MIF1-OE_2

Fig S5

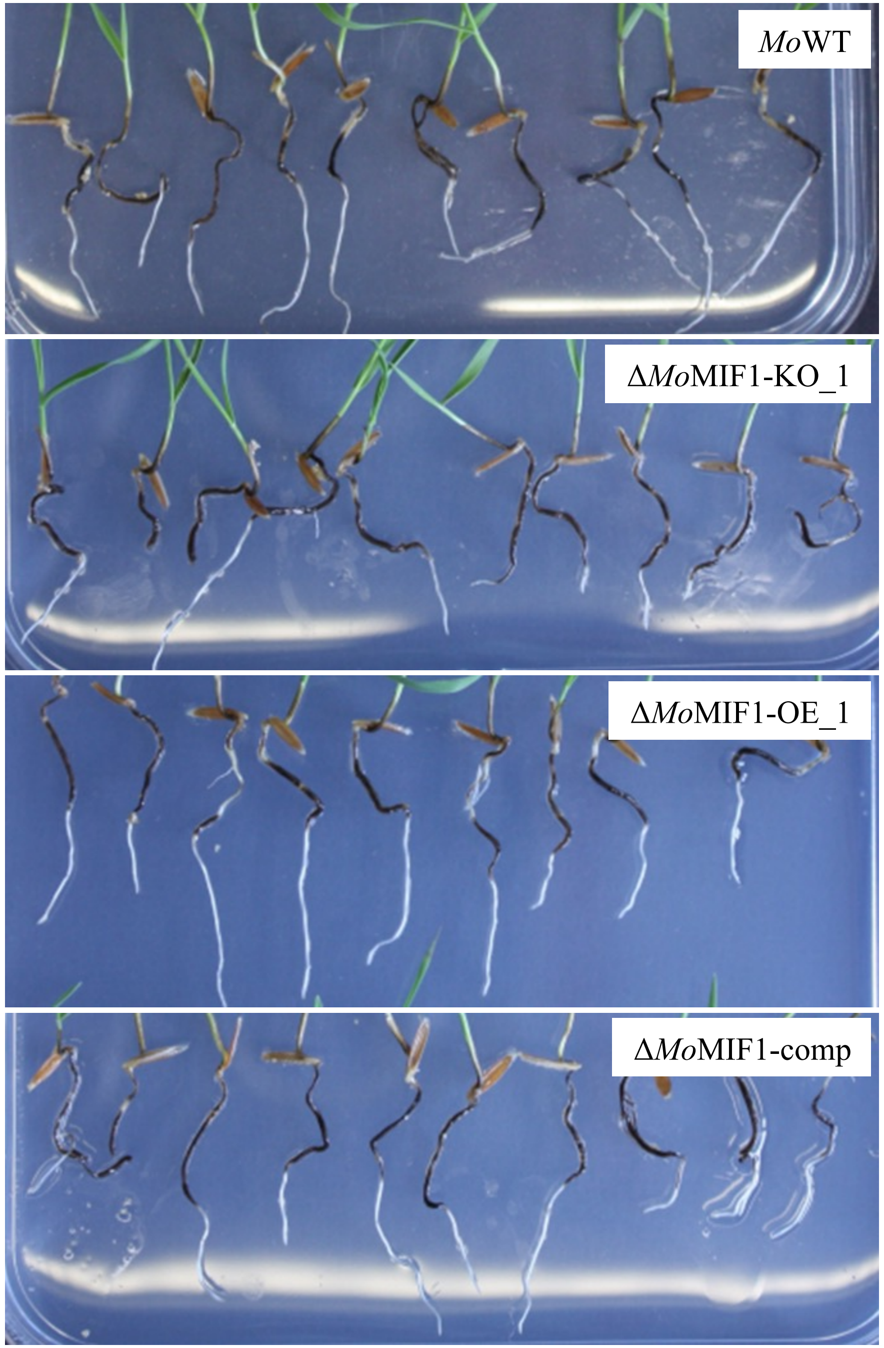

### Fig S6

Sequence: ***Mo*MIF1_MGG_05693T0_protein_coding**

```
                      1          11         21         31         41         51         61         71         81         91
Modifications      1
MoMIF_MGG_056         MAFRPNPVGV GPSIPEEDEE QPQPQPQHLL TGEAGKKDRH PPRSRLQLLA RSSLQRLSFA HRKADLLQRQ HHQSSSQTHR QHNKMSDRQS LDSTSDNCAM

Modifications    101                                                            O
MoMIF_MGG_056         REIERGPPGD RKNNARSRMS QVDVLASKRR STFFEDTFAA SKGEKSGAKA ATAMTERIRS EAIVMAEVKT NVILSDEFTF ITELSYNLSL RYQRPVSSIV

Modifications    201                                              O  O O
MoMIF_MGG_056         VSVQHGACMM YGGTFEPAYS MTIFALPSQM RPTTNKRNAV MIQMHMDEVL GVPSSRGIVR FVPMPEDNVA VSGRTIGSEI TELAREAGID LIDDDEGETL

Modifications    301
MoMIF_MGG_056         SKRRSVKLKN RLSVRKSFSN FKDHSRSGSR ELTPPLPSPP LEPGYSHDGV RERLGPGPND PMPVRKTRRR KSFVASIFGW PREDTNRPPI PL
```

Identified *Mo*MIF1 peptides in sterile intercellular washing fluid (IWF) of Bd21-3 infected with Δ*Mo*MIF1-OE_1 are highlighted in green

Fig S7a

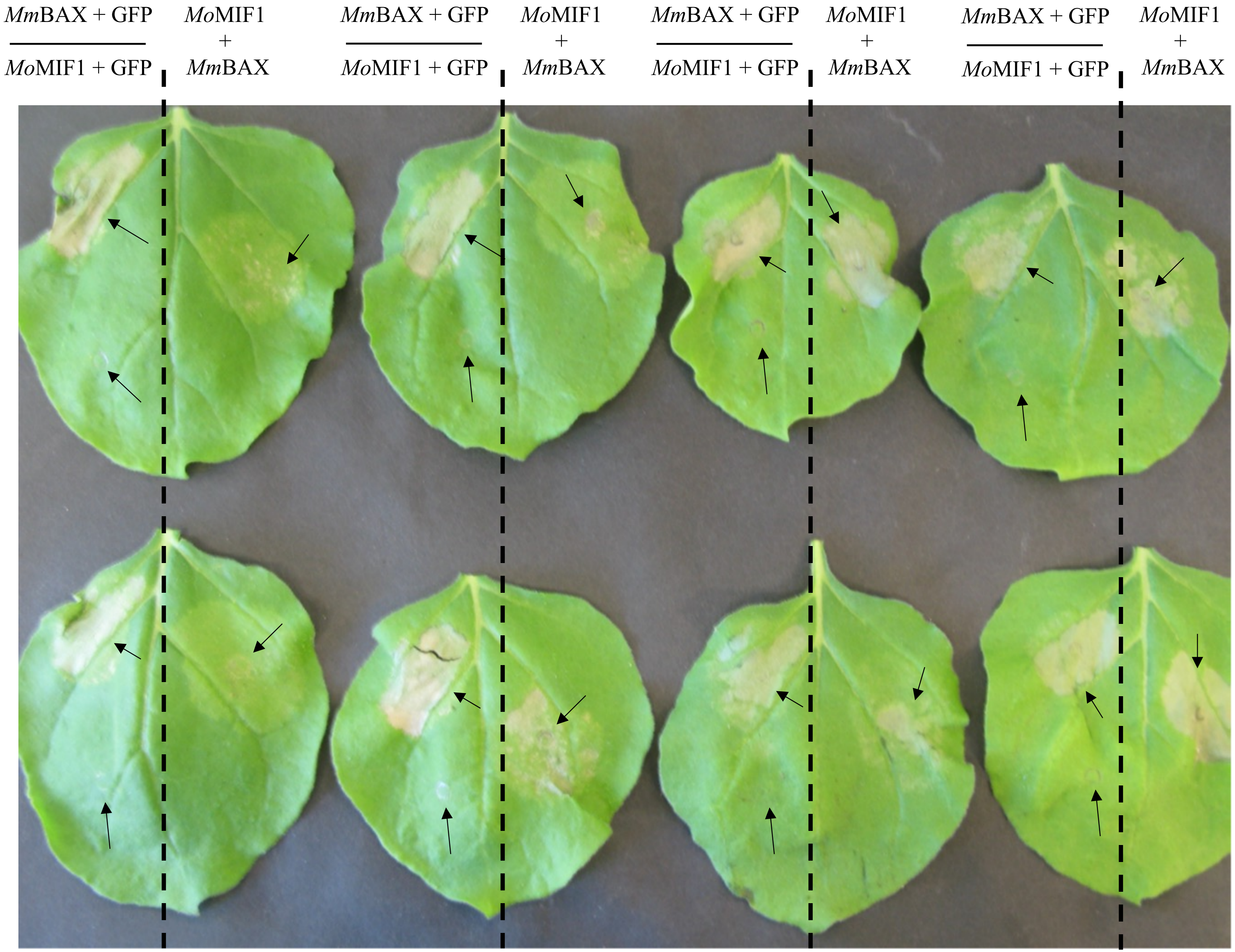

MmBAX + GFP / MoMIF1 + GFP
MoMIF1 + MmBAX
MmBAX + GFP / MoMIF1 + GFP
MoMIF1 + MmBAX
MmBAX + GFP / MoMIF1 + GFP
MoMIF1 + MmBAX
MmBAX + GFP / MoMIF1 + GFP
MoMIF1 + MmBAX

Fig S7b

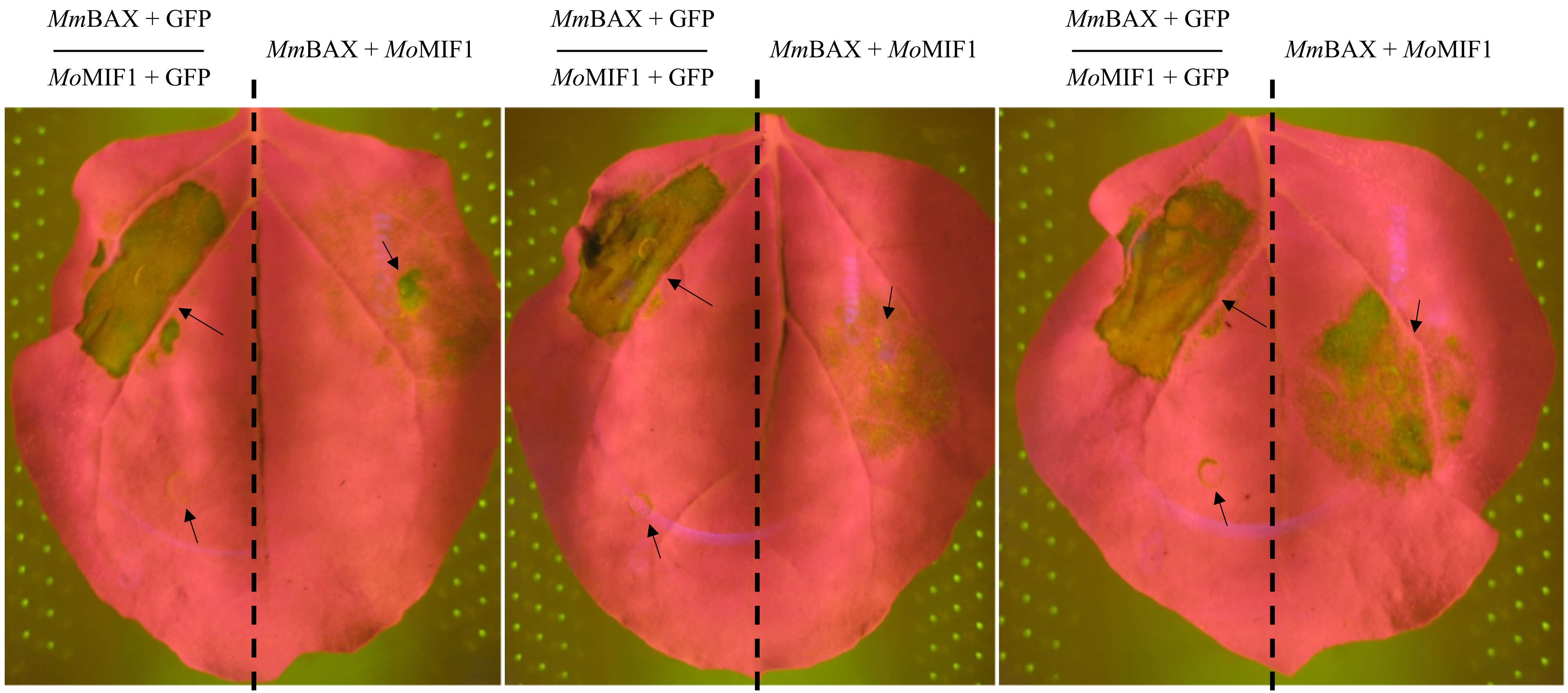

*Mm*BAX + GFP
*Mo*MIF1 + GFP
*Mm*BAX + *Mo*MIF1
*Mm*BAX + GFP
*Mo*MIF1 + GFP
*Mm*BAX + *Mo*MIF1
*Mm*BAX + GFP
*Mo*MIF1 + GFP
*Mm*BAX + *Mo*MIF1

Fig S7c

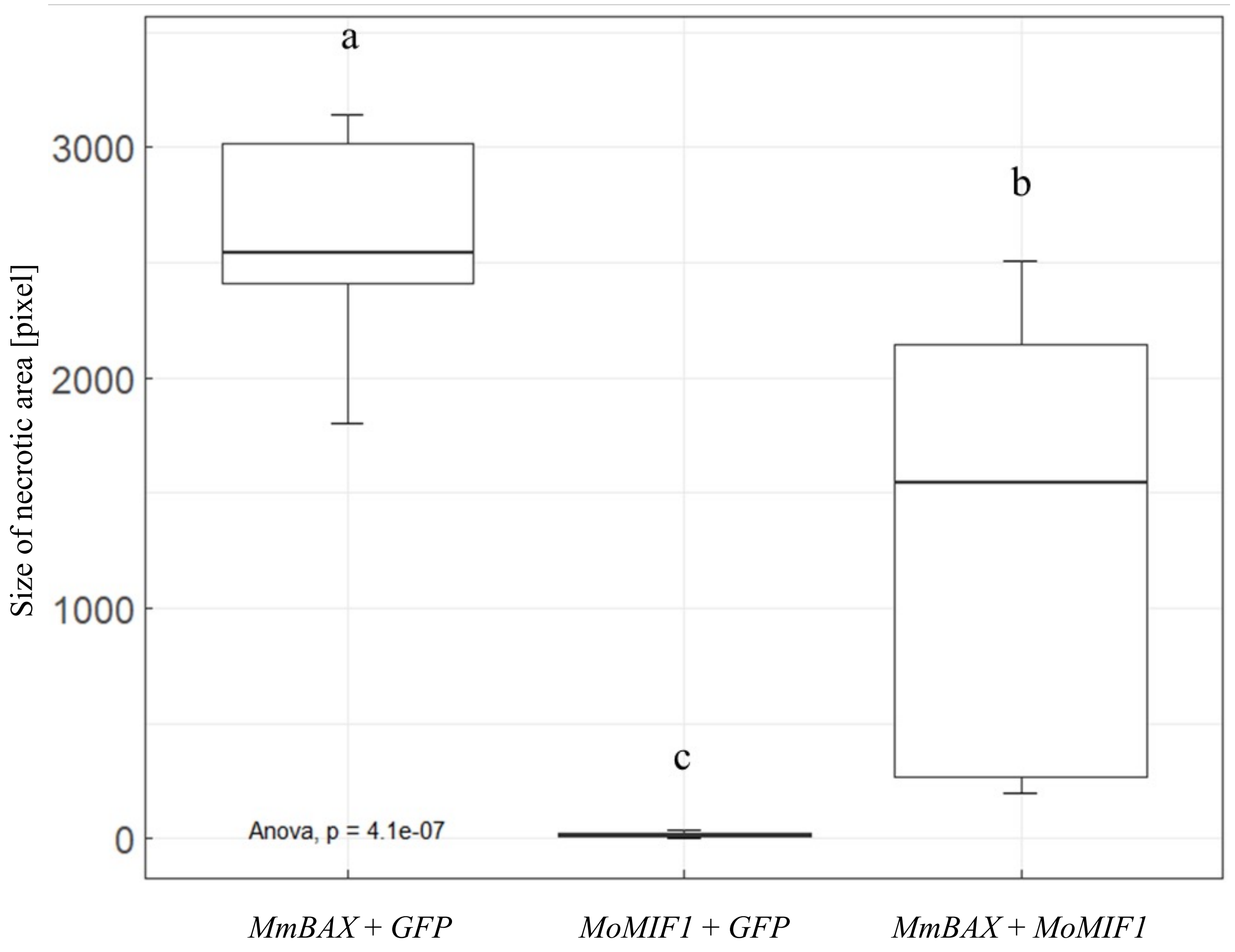

a
b
c
3000
2000
1000
0
Size of necrotic area [pixel]
Anova, p = 4.1e-07
MmBAX + GFP
MoMIF1 + GFP
MmBAX + MoMIF1
