## supplement table for "A key regulator with a MIF-like domain orchestrates cellular differentiation and virulence in the fungal pathogen *Magnaporthe oryzae*"

Table S1

| **Primer Name** | **Sequence 5’ 🡪 3’** | **Target; Use** |
| --- | --- | --- |
| qMoActin_R | ATCTTCTCTCGGTTGGACTTGG | *Mo Actin*; qPCR expression |
| qMoActin_F | AGCGTGGTATCCTCACTTTGC |  |
| qMoGPD_F | CCTCGTCCATCTTCGACGC | *Mo Glyceraldehyde-3-phosphate dehydrogenase*; qPCR expression |
| qMoGPD_R | GACACGACGGCTGTAGCC |  |
| qBdUbi10_R | TGGTACACAGGCATAACACTGACG | *Bd Ubiquitin* 10; qPCR expression |
| qBdUbi10_F | TGGACTTGCTTCTGTCTGGGTTCA |  |
| qMoMIF1_R | TGCCAGGGCAGTGATCTCCTTC | *Mo Macrophage migration inhibitory factor 1*; qPCR expression |
| qMoMIF1_F | GGTCTTGAGAGCGGTGGTATCCATG |  |
| sj-1249 | tggtggcaggatatattgtggtgtaaacaaGGTTTGTCTAGCTTGTCCGTTCCTC | *Cloning for ΔMoMIF1-KO_1* |
| sj-1250 | tcagttaacgtcgacATAGACGTTGCAGAGAGCTACGAGC | *Cloning for ΔMoMIF1-KO_1* |
| sj-1251 | ctctgcaacgtctatGTCGACGTTAACTGATATTGAAGGAGCATTTTTTG | *Cloning for ΔMoMIF1-KO_1/2* |
| sj-1252 | atttaaaaaaaatttGTCGACGTTAACTGGTTCCCGGTCG | *Cloning for ΔMoMIF1-KO_1/2* |
| sj-1253 | ccagttaacgtcgacAAATTTTTTTTAAATTTTTTTTATTTAGATCC | *Cloning for ΔMoMIF1-KO_2* |
| sj-1254 | aaacgctcttttctcttaggtttacctgcaCCAAGGCAACAAGGTAACTAAAATC | *Cloning for ΔMoMIF1-KO_2* |
| sj-1626 | ggctggtggcaggatatattgtggtgtaaacaaCGGCTTTAATCTTGCTTTAACCAGG | *Cloning for ΔMoMIF1+GFP* |
| sj-1627 | tttttttctcgctctcagAGCTTTTCGACCTGAATTGTTGGTAC | *Cloning for ΔMoMIF1+GFP* |
| sj-1628 | tcaggtcgaaaagctCTGAGAGCGAGAAAAAAAAACTCTTCGTCG | *Cloning for ΔMoMIF1+GFP* |
| sj-1629 | ttgttttcatggatcCTAAATCTCGGTGACGGGCAGGACC | *Cloning for ΔMoMIF1+GFP* |
| sj-1630 | gtcaccgagatttagGATCCATGAAAACAAGCATTTGAACACACGGGGTGG | *Cloning for ΔMoMIF1+GFP* |
| sj-1631 | aaacgctcttttctcttaggtttacctgcaTTGGGTAGTCGGCGGCGCGTTGTCT | *Cloning for ΔMoMIF1+GFP* |
| sj-1632 | attatcatcaaaaagagagcaccaaaccgccaccATGGCTTTTCGACCAAACCCTGTTGG | *Cloning for ΔMoMIF1+GFP* |
| sj-1633 | cccggtgaacagctcctcgcccttgctcacCAGCGGTATCGGTGGTCGGTTGGTG | *Cloning for ΔMoMIF1+GFP* |
| sj-1634 | attatcatcaaaaagagagcaccaaaccgccacATGGCTTTTCGACCAAACCCTGTTG | Cloning for ΔMoMIF1-OE_1 |
| sj-1635 | acgatctgcagccgggcggccgctttacttTTACAGCGGTATCGGTGGTCGGTTG | Cloning for ΔMoMIF1-OE_1 |
| sj-1636 | agaaaaaaaaactcttcgtcggattatcatgccgcaATAAATGTAGGTATTACCTGTACATTTTATTTATTC | Cloning for ΔMoMIF1-OE_2 |
| sj-1637 | tggtcgaaaagccatTTTGAAGATTGGGTTCCTACGAAAG | *Cloning for ΔMoMIF1-OE_2* |
| sj-1638 | aacccaatcttcaaaATGGCTTTTCGACCAAACCCTGTTG | *Cloning for ΔMoMIF1-OE_2* |
| sj-1639 | gccaaatgtttgaacgatctgcagccgggcTTACAGCGGTATCGGTGGTCGGTTG | *Cloning for ΔMoMIF1-OE_2* |
| sj-1642 | gataccgctgtaaGATCGTTCAAACATTTGGCAATAAAGTTTCTTAAGATTGAATCC | *Cloning for ΔMoMIF1-comp* |
| sj-1643 | gttttcatggatcCTAAATCTCGGTGACGGGCAGGACC | *Cloning for ΔMoMIF1-comp* |
| sj-1708 | ctggctggtggcaggatatattgtggtgtaaacaaAATGACGACAAGGAAGAGGGACGAAGAAAATTTAACAAAG | *Cloning for ΔMoMIF1-comp* |
| sj-1709 | ctcctcgcccttgctcacCAGCGGTATCGGTGGTCGGTTGGTG | *Cloning for ΔMoMIF1-comp* |
| sj-1726 | cccgtcaccgagatttagGATCCATGAAAACAAGCATTTGAACACACGGGGTGG | *Cloning for ΔMoMIF1-comp* |
| sj-1727 | ctaataaacgctcttttctcttaggtttacctgcaTTGGGTAGTCGGCGGCGCGTTGTCT | *Cloning for ΔMoMIF1-comp* |
